# Intraflagellar transport protein IFT172 contains a C-terminal ubiquitin-binding U-box-like domain involved in ciliary signaling

**DOI:** 10.1101/2024.11.05.620812

**Authors:** Nevin K. Zacharia, Stefanie Kuhns, Niels Boegholm, Anni Christensen, Jiaolong Wang, Narcis A. Petriman, Anna Lorentzen, Jindriska L. Fialova, Lucie Menguy, Sophie Saunier, Soren T. Christensen, Jens S. Andersen, Sagar Bhogaraju, Esben Lorentzen

## Abstract

Intraflagellar transport (IFT) is a fundamental process driving ciliogenesis in most eukaryotic organisms. IFT172, the largest protein of the IFT complex, plays a crucial role in cilium formation and several disease-causing IFT172 variants have been identified in ciliopathy patients. While IFT172 is tethered to the IFT-B complex via its N-terminal domains, the function of its C-terminal domains has remained elusive. Here, we reveal that the C-terminal part of IFT172 interacts with IFT-A complex subunits, providing a molecular basis for the role of IFT172 in bridging IFT-A and IFT-B complexes. We determine the crystal structure of the C-terminal part of IFT172, uncovering a conserved U-box-like domain often found in E3 ubiquitin ligases. This domain exhibits ubiquitin-binding properties and IFT172 undergoes ubiquitin conjugation in vitro, an activity which is reduced in the C1727R patient ciliopathy variant. We use CRISPR-engineered RPE-1 cells to demonstrate that the U-box-like domain is essential for IFT172 protein stability and proper cilium formation. Notably, RPE-1 cells with heterozygous deletion of the U-box domain show altered TGF-β signaling responses, particularly in SMAD2 phosphorylation levels and AKT activation. Our findings suggest that IFT172, beyond its structural role in bridging IFT-A and IFT-B complexes within IFT trains, harbors a conserved U-box-like domain with potential involvement in ciliary ubiquitination processes and signaling, providing new insights into the molecular mechanisms underlying IFT172-related ciliopathies.

## Introduction

Cilia are hair-like microtubule (MT) based organelles that protrude from the cell surface of vertebrate cells and in several eukaryotic single-celled organisms. Cilia play indispensable roles in cellular motility, extracellular fluid flow, sensing extracellular cues, and coordinating cellular signaling pathways^1^. They are highly conserved from unicellular eukaryotes to humans and can vary greatly in length, copy number, and function. For example, in the unicellular algae *Chlamydomonas reinhardtii* (Cr), beating motions generated by motile cilia propel the cells through media^2^. Additionally, most vertebrate cells possess a single copy of non-motile primary cilia that performs sensory and signaling functions. The sensory capabilities of the primary cilium are attributed to ciliary coordination of signaling pathways such as transforming growth factor β (TGF-β)/bone morphogenetic protein (BMP), platelet-derived growth factor (PDGF), and Hedgehog to drive organismal development and tissue homeostasis^3^. Aberrations to cilium assembly or function leads to a class of human genetic disorders termed ciliopathies^4^. Ciliopathies affect a wide range of organs including the liver, kidney, and skeletal system and often manifest as syndromes such as Bardet Biedl syndrome and Meckel Gruber syndrome with overlapping phenotypes^5^.

The proper assembly and function of cilia require a bidirectional trafficking system known as intraflagellar transport (IFT)^6–8^ to shuttle key ciliary components^9,10^, including membrane-bound receptors such as G protein-coupled receptors (GPCRs) and ion channels that mediate signaling and sensory responses, between the cilium organelle and the cytoplasm^10–12^. IFT is carried out by the 23 subunit IFT complex that polymerizes into IFT trains^8,13^ and traverses along the ciliary MT axoneme. IFT trains assemble at the ciliary base^14^, associate with ciliary cargo^10,15^ and undergo anterograde transport from the ciliary base to the tip powered by the Kinesin-2 motor^8,16^. A major remodeling of the anterograde IFT trains occurs at the ciliary tip^17^, forming retrograde IFT trains that have a distinct ultrastructure^18^, which subsequently traverse from the tip to the ciliary base powered by the Dynein-2 motor^19,20^. The IFT complex associates with the octameric BBSome complex to facilitate the targeted ciliary exit of several proteins^21,22^. The IFT trains are primarily understood to function as adaptors between the molecular motors and ciliary cargo to facilitate the dynamic localization of signaling components.

Ubiquitination is a well-studied posttranslational modification^23^ that was recently shown to be crucial in ciliary homeostasis^24–26^. Ubiquitination is a multi-step enzymatic cascade where a ubiquitin molecule is shuttled from an E1 enzyme to the E2 enzyme and finally transferred to the substrate protein mediated by an E3 ligase^23^. Ubiquitination is known to be important in flagella disassembly in *Cr*^27^. Upon initiation of flagellar resorption, there is an upregulation of ubiquitination events, specifically that of tubulin subunits, facilitating ciliary disassembly^27,28^. Ubiquitination is also critical to ciliary signaling. The ciliary Hedgehog signaling pathway is a well-studied example of how regulatory ubiquitination events control signal dependent repression and activation of ciliary signaling^25,26,29,30^. In the absence of the Sonic Hedgehog ligand (SHH), the ciliary levels of the signal-transducing protein smoothened (SMO) is kept low for pathway repression^29^. This is achieved by the ubiquitination of SMO by the ciliary localized ubiquitin ligase WWP1, an event that subsequently targets SMO for IFT dependent ciliary exit^25,29^. Upon SHH binding, the ciliary exit of the ligand bound PTCH1 receptor and negative pathway regulator GPR161 occurs by the targeted ubiquitination of these receptors^26,30^, leading to pathway activation. In the case of several GPCRs, such as GPR161 and SSTR3, signal dependent ciliary exit specifically relies on the ubiquitination of these receptors and subsequent IFT dependent ciliary exit^26^. *Bona fide* ubiquitin recognition domains are yet to be identified in the IFT complex. Instead, the BBSome complex was shown to facilitate the recognition of these ubiquitinated GPCRs mediated by the TOM1L2 adaptor protein that harbors VHS and GAT domains capable of ubiquitin binding^31^. Moreover, a recent analysis of ubiquitinated proteins in the cilium of RPE1 cells identified signaling-related proteins to be enriched in the data set^32^. Specifically, components of the Wnt, Notch, and TGF-β pathway were identified as targets of ciliary ubiquitination events. The BBSome complex, which mediates signal dependent ciliary exit of a wide array of ubiquitinated GPCRs, is in turn dependent on ubiquitination for proper assembly and function^33^. The cilium associated ubiquitin ligase PJA2 ubiquitinates the BBSome subunits BBS1 and BBS4, which is essential for normal BBSome assembly and ciliary GPCR trafficking^33^. Cilium associated ubiquitin ligases and ubiquitin binding domains thus emerge as crucial players in maintaining ciliary homeostasis.

IFT172 is the largest subunit in the IFT complex and is part of the IFT-B2 subcomplex^34^. The WD40 repeat containing N-terminal domain of the protein mediates interaction with the subunits IFT57 and IFT80, thereby maintaining IFT172 association with the IFT-B complex^34,35^. The C-terminal tetratricopeptide repeats (TPRs) of IFT172 remains rather uncharacterized in terms of structure and function despite the fact that IFT172 C-terminal variants have been detected in a cohort of ciliopathy patients with severe skeletal ciliopathy syndromes (Jeune syndrome and Mainzer-Saldino syndrome)^36^. Another cohort of patients with phenotypically slightly different ciliopathy features resembling BBS also presented with IFT172 variations predominantly resulting in retinal defects^37^. Lastly, an isolated case of a patient exhibiting growth retardation was reported to have compound heterozygous IFT172 C-terminal mutations^38^. The importance of the IFT172 C-terminus is further exemplified by prior studies on various *in vivo* models. IFT172^wim^ mice carry a C-terminal point mutation (L1564P) causing major developmental defects and embryonic lethality caused by a complete loss of cilium in the embryonic node^39^. The *Chlamydomonas reinhardtii* strain *fla11* carry the point mutation L1615P at the C-terminal TPRs of IFT172 resulting in flagellar resorption and IFT particle accumulation at the ciliary tip of green algae^40^, a phenotype that was recapitulated in *Tetrahymena thermophila* models having progressive IFT172 C-terminal deletions^41^. Furthermore, it was observed that the *Chlamydomonas reinhardtii fla11* strain accumulates ubiquitinated proteins in the cilium^42^, which suggests that IFT172 could be involved in ciliary ubiquitination events. However, the molecular bases for these observations are currently not known.

In this study, we focused on structural and functional characterization of the IFT172 C-terminal region to reveal an interaction with IFT-A components and a C-terminal ubiquitin-binding U-box domain that may be involved in ciliary ubiquitination and signaling pathways.

## Results

### IFT172_968-C_ interacts directly with subunits of the IFT-A complex

While the N-terminal β-propeller of CrIFT172 interacts with CrIFT57 and a central region (residues 626-785) interacts with CrIFT80, little is known about the function of the C-terminal 800 residues of IFT172. This region is predicted to fold into 20 TPRs followed by a small 70-80 residue domain of unknown function (Fig. 1A).

**Figure 1.**
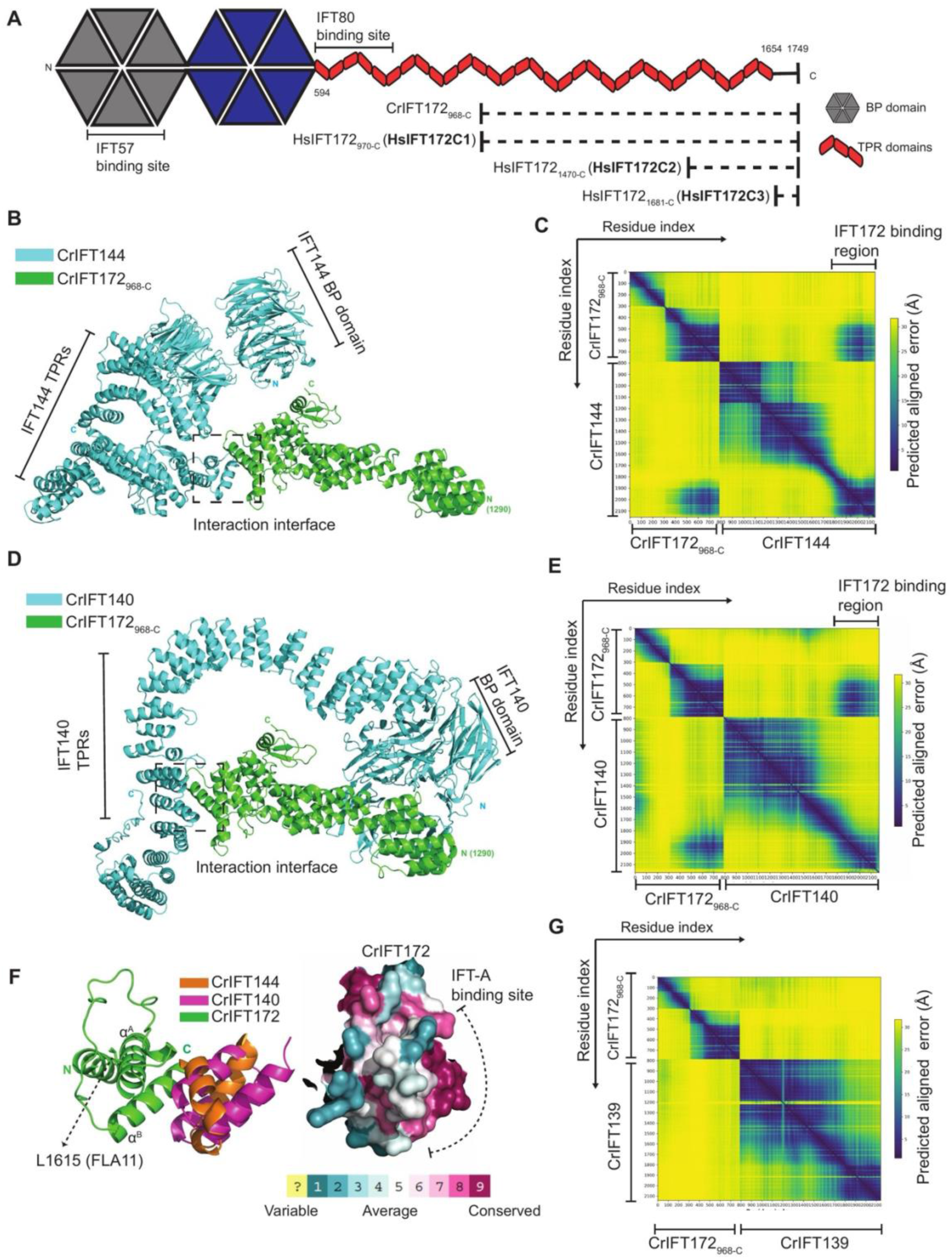
The C-terminal TPRs of IFT172 constitute a binding site for IFT-A subunits. **(A)** Schematic representation of Homo sapiens (Hs) IFT172 protein domain organization. β-propeller and TPR (tetratricopeptide repeat) domains are indicated. Solid lines represent previously identified IFT57 and IFT80 binding sites. Dashed lines show various truncated constructs of *Chlamydomonas reinhardtii* (Cr) IFT172 and HsIFT172 used in this study. **(B)** Cartoon representation of the AlphaFold predicted structural model for a complex between CrIFT172_968-C_ and CrIFT144. The interaction interface is highlighted. **(C)** Predicted Aligned Error (PAE) plot for the AlphaFold model in panel B. X- and Y-axes show indexed residues from each protein used for PAE calculation. Low PAE scores indicate high confidence in the predicted interaction interface. **(D)** Cartoon representation of the AlphaFold predicted structural model for a complex between CrIFT172_968-C_ and CrIFT140, highlighting the interaction interface. **(E)** PAE plot for the AlphaFold model in panel D, showing high confidence (low PAE values) for the interaction interface residue pairs. **(F)** Left: Structural superposition of interaction interfaces from AlphaFold models in panels B and D. Subunits are colored as indicated. IFT144 and IFT140 are predicted to associate with an identical binding site formed by IFT172 TPR helices αA and αB. The L1615P temperature-sensitive mutation identified in the *fla11* strain of *C. reinhardtii* maps onto helix αA. Right: Surface amino acid conservation map for the IFT144/140 binding site in IFT172, color-coded as indicated. **(G)** PAE plot for an AlphaFold predicted structure of the CrIFT172_968-C_-CrIFT139 complex. High PAE scores for all inter-chain residue pairs suggest that IFT172 and IFT139 do not interact directly.

We purified the C-terminal fragment of IFT172 protein (His-tagged CrIFT172_968-C_) to homogeneity (Fig. S1A) and carried out pull-down assays from an extract of isolated *Chlamydomonas reinhardtii* cilia (Fig. S1B). His-tagged Tobacco Etch Virus (TEV) protein was used as a negative control when detecting CrIFT172_968-C_ interactors by mass spectrometry (MS) (Figs. S1C-D). Based on the resulting volcano plot (Fig. S1C, see also M&M), 10 proteins were annotated as interactors of CrIFT172_968-C_ (Fig. S1D). Consistent with the fact that the CrIFT172_968-C_ construct does not contain the IFT-B complex associating domains, no IFT-B proteins were found as interactors (Fig. S1D). Interestingly, however, several components of the IFT-A complex were identified as interaction partners of CrIFT172_968-C_ including IFT144, IFT140, and IFT139. Furthermore, a CH-domain containing protein of the central apparatus of the cilium as well as six additional proteins without a published ciliary localization were identified as IFT172 interactors (Fig. S1D).

To assess which of the 10 significant hits are direct interaction partners of CrIFT172_968-C_, structural modeling using AlphaFold multimer (AF-M)^43,44^ was carried out. This revealed 3 high-confidence hetero-dimeric complexes between CrIFT172_968-C_ and each of the proteins: IFT144, IFT140, and a UBX-domain containing protein (uniprot: A0A2K3DQG3, CHLRE_06g293900v5) (Fig. 1B-E and S1E-F). None of the other seven candidate proteins, including IFT139 (Fig. 1G), was predicted with any confidence to interact directly with CrIFT172_968-C_, and may constitute indirect interactors.

The structural model of the CrIFT172-IFT144 complex shows that the interaction is formed by the most C-terminal TPR of IFT172 (residues 1604-1669) and TPR helices of IFT144 (residues 1220-1250, see Fig. 1B). Surprisingly, the interaction with IFT140 (residues 1125-1162) utilizes the same TPR helices of CrIFT172 as the IFT144 interaction (Fig. 1D and 1F). The structural models suggest that the interactions of IFT172 with IFT140 and IFT144 are mutually exclusive (Fig. 1F). Both IFT172-IFT144 and IFT172-IFT140 interactions are predicted with high confidence according to the Predicted Aligned Error (PAE) plots (Figs. 1C and 1E). The conservation of the IFT140 and IFT144 interaction surface on IFT172 across ciliated organisms suggests a conserved function in binding IFT-A subunits (Fig. 1F).

We note that the IFT172 interacting helices of IFT144 are surface exposed in the cryo-EM structures of complete IFT-A complexes^45^ and thus free to interact with IFT172 in IFT trains. Indeed, in the published cryo-ET structures of anterograde IFT-trains in Cr, the IFT172-IFT144 complex described here is observed^46^. Interestingly, cryo-ET structures of retrograde IFT trains demonstrate significant rearrangements of IFT-A and B sub-complexes^47^ and reveal the same IFT172-IFT140 interaction as shown in Fig. 1D-F. The mutually exclusive interactions between IFT172 C-terminus and IFT140 or IFT144 thus underpin the different architectures of anterograde and retrograde IFT trains. Fig. 1F highlights the L1615 residue, which is mutated to a proline in the *fla11* strain resulting in a retrograde IFT defect phenotype in Cr^40^ and accumulation of ubiquitinated proteins^42^. Interestingly, the L1615 residue is located in one of the two helices of IFT172 that interact directly with IFT140 or IFT144 (annotated as helix αA in Fig. 1F). This L1615P mutation will disrupt the helix αA, likely resulting in impaired binding of IFT172 to the IFT-A complex. This suggests that the molecular basis for the ciliary defects observed in *fla11* is caused by impaired IFT172-IFT-A association.

The uncharacterized UBX-domain-containing protein was predicted by AF-M to be a potential direct IFT172 interactor (Fig. S1E-F). UBX-domain-containing proteins generally function as cofactors that link the proteasome to ubiquitinated substrates, playing crucial roles in protein degradation and cellular quality control mechanisms^48^. They often act as adaptors, facilitating recognition and processing of ubiquitinated proteins by proteasomes. The identification of a UBX-domain protein as an IFT172 interactor is consistent with previous findings demonstrating that the UBX-domain protein UBXN10 directly interacts with the IFT-B complex via CLUAP1/IFT38, recruiting the VCP/p97 ATPase to regulate IFT-B complex integrity and ciliogenesis^94^. This suggests that UBX-domain proteins may serve broader roles as regulators of IFT complex remodeling. However, despite our interest in the interaction between IFT172 and UBX-domain-containing proteins, we could not recombinantly express the UBX-domain-containing protein in a soluble form, which precluded further experimental characterization of its relationship with IFT172.

### The crystal structure of human IFT172 reveals a C-terminal U-box-like domain

To gain deeper insights into the molecular architecture and potential functional domains of IFT172, we pursued crystallographic studies of IFT172C. Structural models of CrIFT172 predicted a small domain at the C-terminus that does not form TPRs (Fig. 2A). This domain, which contains mixed α/β secondary structures, is linked to the IFT144 and IFT140 binding helices of IFT172 through a loop region (denoted as L0) that spans 30 residues. This arrangement places the C-terminal domain roughly 25Å away from the IFT144 and IFT140 binding helices of IFT172 (Fig. 2A). In our pursuit of experimental validation, we isolated and crystallized a C-terminal construct of *Homo sapiens* (Hs) IFT172_1470-C_ (hereafter referred to as HsIFT172C2, see Fig. S2A for purification). This construct includes the IFT144 and IFT140 binding site and the small non-TPR C-terminal domain identified in the AlphaFold model of CrIFT172 (Fig. 2A). X-ray diffraction data were collected to 2.1Å resolution from native crystals and 2.8Å resolution at the peak wavelength of selenomethionine-substituted crystals (Table 1). Crystallographic phase information was obtained by combining molecular replacement (AlphaFold model of HsIFT172C2) and single-wavelength anomalous dispersion yielding an electron density map of excellent quality (Fig. S2B). This map allowed for the modeling of the entire structure, excluding a small loop (residues 1656-1657) and the seven most C-terminal residues. The crystal structure reveals that HsIFT172 contains a small domain (residues 1683-1749, referred to as HsIFT172C3) at the C-terminus, which consists of a small 2-stranded β-sheet followed by two α-helices (see Fig. 2B). The preceding TPRs and connecting loop (loop L0, colored magenta in Fig. 2B) pack against this C-terminal domain via a central hydrophobic core and several peripheral polar interactions (Fig. S2E). Interestingly, numerous missense variants from ciliopathy patients^36–38^ map to this interface and are highlighted in Fig. 2B as yellow sticks.

**Figure 2.**
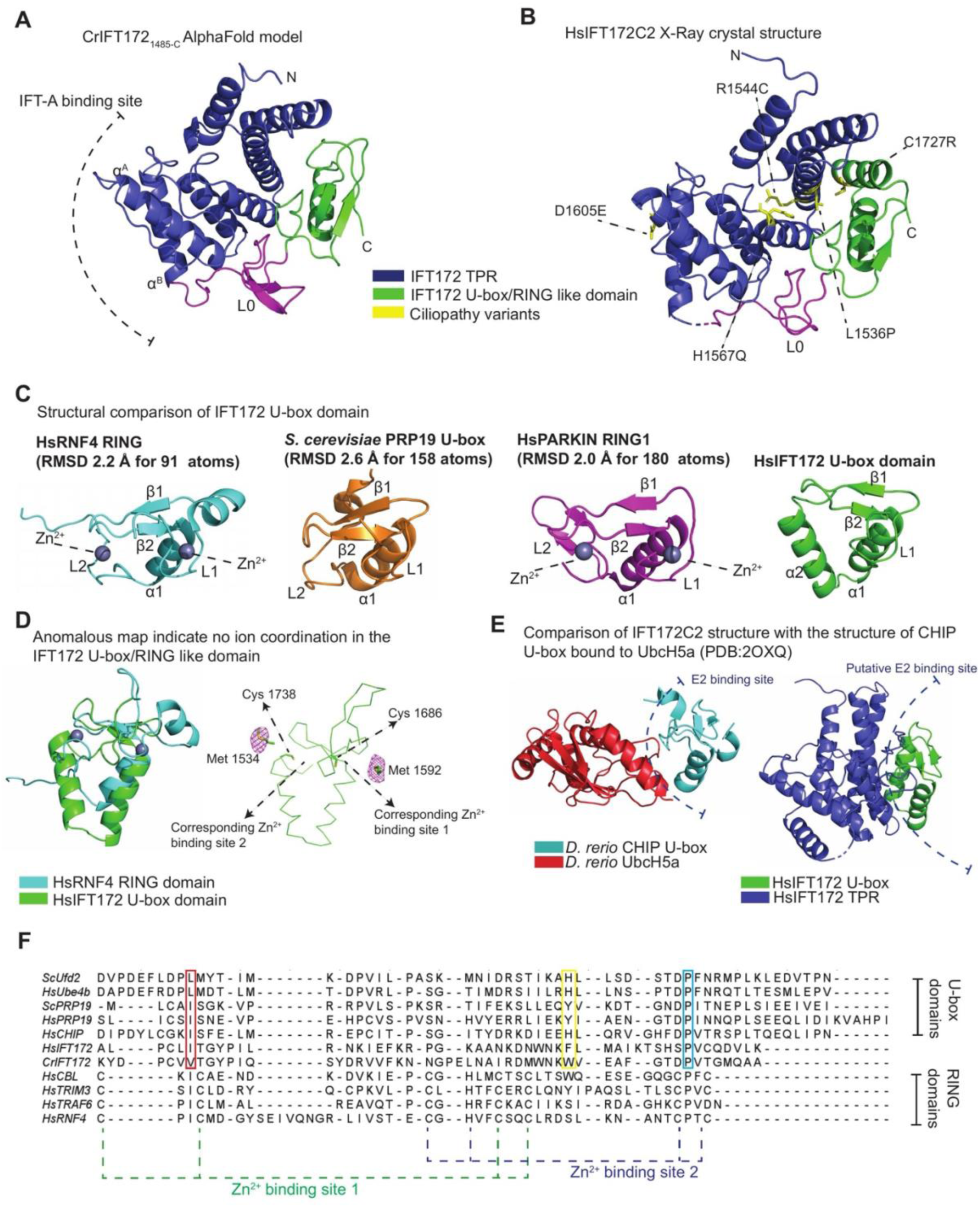
IFT172 contains a U-box like domain distal to the IFT-A binding site. **(A)** Cartoon representation of the AlphaFold-predicted structure of the C-terminal residues 1485-1755 of CrIFT172. This globular domain comprises several TPR helices, including the IFT140/144 binding αA and αB helices, followed by a loop region (L0) and a U-box/RING-like motif. **(B)** Cartoon representation of the 2.1Å resolution X-ray crystal structure of HsIFT172C2. The structure exhibits a similar fold to the CrIFT172_1485-1755_ AlphaFold model, composed of TPR helices followed by a loop region (L0) and a U-box/RING-like motif. **(C)** Structural comparison of the U-box/RING-like motif in HsIFT172 with canonical RING domains of HsRNF4 (PDB: 4PPE), RING1 domain of PARKIN (PDB: 6HUE), and U-box domain of ScPRP19 (PDB: 6BAY), indicating the corresponding RMSD after superposition with the IFT172 motif. The U-box/RING-like motif in HsIFT172 superposes well with several structural components of U-box/RING domains, including the first loop (L1), the following beta strands (β1 and β2), and the helix (α1). A major difference in this HsIFT172 motif is the replacement of the characteristic second loop region (L2) found in U-box/RING domains with an alpha helix (α2). **(D)** (Left) Structural superposition of the IFT172 U-box/RING-like domain with the RING domain of RNF4, identifying corresponding Zn^2+^ binding sites. Zn^2+^ ions in RNF4 are depicted as grey spheres. (Right) Anomalous density map represented as a magenta mesh contoured at 5σ, obtained from HsIFT172C2 selenium-methionine substituted protein crystals. Anomalous density depicted in proximity to the U-box/RING-like domain. Cys residues and neighboring Met residues in the predicted Zn^2+^ binding site of IFT172 are represented as sticks. **(E)** (Left) Structure of the U-box domain in *D. rerio* CHIP bound to *D. rerio* UbcH5a (PDB: 2OXQ), indicating the E2 binding site on the CHIP U-box domain. (Right) Comparison of HsIFT172C2 crystal structure with PDB: 2OXQ indicates the putative E2 binding site on IFT172 U-box is occluded by the IFT172 TPR domain. **(F)** Sequence alignment of U-box domains in HsIFT172 and CrIFT172 with several canonical U-box/RING domains. Zn^2+^ coordinating residues on RING domains are highlighted. Selected functionally relevant residues on U-box domains are depicted in boxes, and corresponding residues are also observed in the IFT172 U-box domain sequence.

**Table 1:** X-ray diffraction data collection and refinement statistics (PDB: 9H2D).

| Two protein copies / asymmetric unit | HsIFT172<sub>1470-C</sub><br>Native | *HsIFT172*<sub>1470-C</sub><br>Selenium methionine |
| --- | --- | --- |
| Data collection |  |  |
| Wavelength (Å) | 1.006 | 0.9785 |
| Resolution range (Å) | 45.25-2.07<br>(2.10-2.07) | 47.02-2.78<br>(2.95-2.78) |
| Space group | P 1 2<sub>1</sub> 1 | P 1 2<sub>1</sub> 1 |
| Unit cell (Å, °) | a = 46.97<br>b = 90.51<br>c = 67.42<br>$\alpha$ = 90<br>$\beta$ = 97.42<br>$\gamma$ = 90 | a = 47.47<br>b = 90.63<br>c = 67.81<br>$\alpha$ = 90<br>$\beta$ = 97.93<br>$\gamma$ = 90 |
| Total reflections | 225982 (10379) | 84456 (12152) |
| Unique reflections | 34028 (1631) | 26158 (3916) |
| Multiplicity | 6.6 (6.4) | 3.2 (3.1) |
| Completeness (%) | 99.5 (96.2) | 93.0 (85.9) |
| Mean I/sigma | 9.7 (1.3) | 8.3 (0.7) |
| R<sub>p</sub>im | 0.037 (0.483) | 0.077 (1.65) |
| CC1/2 | 0.999 (0.777) | 0.998 (0.494) |
| Refinement |  |  |
| Resolution range (Å) | 45.25-2.10<br>(2.17-2.10) |  |
| Reflections for refinement | 32556 (3050) |  |
| Reflections for R<sub>free</sub> | 1564 (151) |  |
| Protein residues | 532 |  |
| Number of atoms | 4423 |  |
| Protein | 24213 |  |
| Ligands | 31 |  |
| Water (solvent) | 179 |  |
| R-work | 0.189 (0.367) |  |
| R-free | 0.238 (0.370) |  |
| Ramachandran favored (%) | 97.35 |  |
| Ramachandran outliers (%) | 0.00 |  |
| RMS bonds (Å) | 0.01 |  |
| RMS angles (°) | 1.23 |  |
| Average B-factors (Å<sup>2</sup>) | 65.14 |  |

Statistics for the highest resolution shell are shown in parentheses.

Using this C-terminal HsIFT172C3 domain structure in a Distance-matrix ALIgnment (DALI)^49^ search against the Protein Data Bank (PDB), structural similarities to RING domain-containing eukaryotic E3 ubiquitin ligase as well as possible prokaryotic RING domain homologues (Zinc finger proteins) were uncovered (Fig. S2C). RING domains are among the most abundant classes of ubiquitin ligase domain in humans^50^. Therefore, we sought to evaluate the possible structural similarity of HsIFT172C3 to various ubiquitin ligase domains. Fig. 2C presents a structural comparison of the HsIFT172C3 domain with representative domains from different classes of E3 ubiquitin ligases. HsIFT172C3 superposes with RING and U-box domains with root-mean-square deviations (RMSDs) of 2.0-2.6Å with structure-based sequence identities of 14-17% (Fig. 2C and 2F). The common structural architecture of the U-box/RING domain is characterized by an N-terminal loop (L1) followed by a small two-stranded antiparallel β-sheet (β1 and β2), an α-helix (α1) and a C-terminal loop (L2) (Fig. 2C). The RING domain fold is stabilized by the coordination of two Zn^2+^-ions whereas the structurally and functionally similar U-box domain is instead stabilized by a network of polar contacts^51–53^. HsIFT172C3 has a U-box/RING domain structure with the important difference that L2 is replaced by an additional helix α2 (Fig. 2C). Interestingly, the CrIFT172 C-terminal domain is predicted to have the L2 loop and more closely resemble canonical U-box/RING domains (Fig. 2A). Upon inspecting the experimental electron density maps of HsIFT172C2, we did not observe any density at the corresponding Zn^2+^-ion binding sites (see Fig. 2D for an anomalous electron density map). This observation is supported by sequence alignments, which show a lack of conservation of Zn^2+^ ion coordinating Cys and His residues in IFT172 proteins (Fig. 2F).

These observations suggest that IFT172C3 is not a classical RING domain but rather resembles a U-box domain. U-box/RING domains mechanistically function by binding the ubiquitin-loaded E2 enzyme (E2-Ub conjugate) thus restricting the E2-Ub conjugate to conformations that facilitate ubiquitin transfer to the substrate^54–57^. Interestingly, structural comparisons with well-characterized U-box domains indicate that the predicted E2 binding site in HsIFT172C3 is structurally occluded by TPRs preceding the U-box domain (Fig. 2E). Structural occlusion of the E2 binding site is a prominent mechanism of self-regulation seen in E3 ligases^58–61^. We conclude that IFT172 contains a U-box-like domain of unknown function at the C-terminus, which is a hotspot for ciliopathy patient variants.

### IFT172 undergoes ubiquitination in the presence of the Ubch5a E2 enzyme

Given the structural similarity of HsIFT172C3 to U-box domains of well-characterized E3 ubiquitin ligases, we decided to test if IFT172 exhibits ubiquitination activity. To this end, various C-terminal constructs of Cr and Hs IFT172 proteins were tested in *in vitro* ubiquitination assays in the presence of purified MmUbe1 (E1 enzyme), ubiquitin, UbcH5a (E2 enzyme) and ATP (Figs. 3A). Appearance of ubiquitin conjugates was observed in reactions containing CrIFT172_968-C_ and HsIFT172_970-C_ (HsIFT172_970-C_ will be referred to as HsIFT172C1 from hereon). In the presence of ubiquitination pathway components (E1, E2, and ATP), ubiquitin ligases have the characteristic property of self-ubiquitination (auto-ubiquitination) *in vitro*^53,62^. Screening activity for HsIFT172C1 with 11 different E2 enzymes revealed the appearance of ubiquitin conjugates only in the presence of members of the Ubch5 E2 enzyme family (Fig. 3B). The molecular mass of the ubiquitin conjugate is consistent with mono-ubiquitinated HsIFT172C1 used in the assay (Fig. 3A and 3B) and is also stained by anti-His antibodies detecting the His-tag on HsIFT172C1 (Fig. 3C). Polyubiquitination is a hallmark of E3 ubiquitin ligases resulting in an observed smear of ubiquitinated products on ub-antibody stained western blots^53^. A characteristic smear is not observed in the case of IFT172 but rather a strong band corresponding to mono-ubiquitination is observed. Interestingly, higher molecular weight ubiquitinated bands of lower intensity, that migrates close to the prominent mono-ubiquitinated IFT172 are observed in several replicates (Fig. 3A, 3B and 3F). These bands could potentially correspond to additional mono-ubiquitination events on IFT172 or the formation of short poly-ubiquitin chains on IFT172.

**Figure 3.**
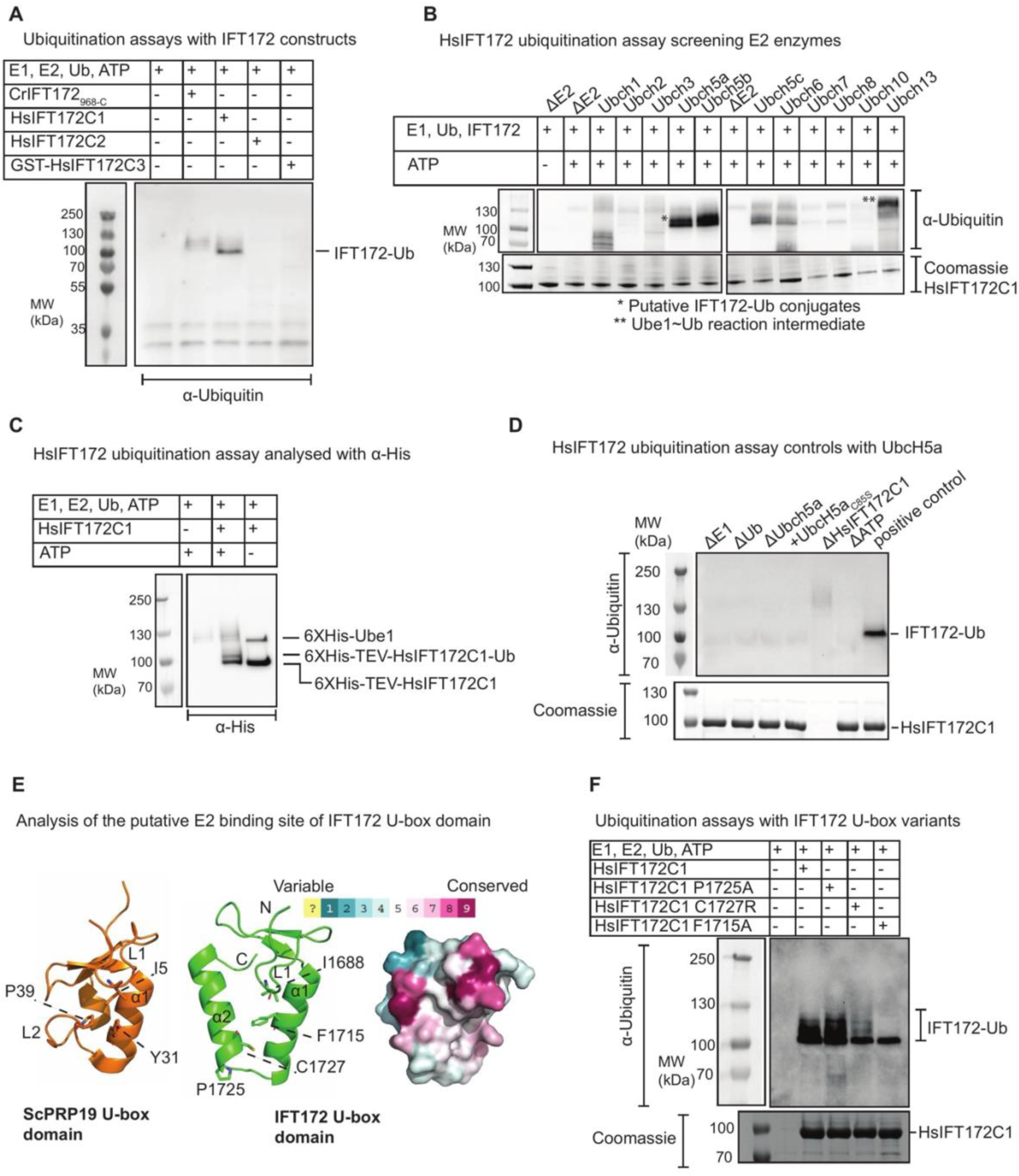
IFT172 exhibits ubiquitin conjugation activity in the presence of UbcH5a. **(A)** Western blot analysis of an *in vitro* ubiquitination assay containing *M. musculus* Ube1(E1), HsUbcH5a (E2), Ubiquitin (Ub) and ATP in the presence of various C-terminal IFT172 constructs as potential E3 ligases. Reactions were visualized by immunostaining with anti-Ubiquitin antibody. **(B)** Western blot analysis of *in vitro* ubiquitination assays containing Ube1 (E1), Ub, ATP and HsIFT172C1, in the presence of 11 different ubiquitin-conjugating E2 enzymes (Abcam #ab139472). Reactions were visualized by immunostaining with (top) anti-Ubiquitin antibody and (bottom) Coomassie staining. Reactions were conducted under non-reducing conditions, accounting for the visualization of significant amounts of E1∼Ub conjugate in the blot. **(C)** *In vitro* ubiquitination assays containing *M. musculus* 6XHis-Ube1, 6XHis-TEV-HsUbcH5a, Ub and 6XHis-TEV-HsIFT172C1. Reactions were visualized by immunostaining with an anti-His tag antibody. **(D)** Western blot analysis of *In vitro* ubiquitination reactions containing *M. Musculus* Ube1(E1), HsUbcH5a (E2) WT/C85S mutant, Ubiquitin (Ub), HsIFT172C1 and ATP. As specified in each reaction, a reaction component was omitted (indicated by Δ) or a mutant component was used instead of the corresponding WT component (indicated by +). Reactions were visualized by immunostaining with (top) anti-Ubiquitin antibody and (bottom) Coomassie staining. **(E)** Analysis of the putative E2 binding site in the (center) HsIFT172 U-box domain and (Left) U-box domain of ScPRP19. Both U-box domains are shown facing the E2 binding site. The PRP19 residues I5, Y31, and P39 whose mutagenesis leads to the loss of its ubiquitin ligase activity are represented as sticks (also highlighted in boxes within the sequence alignment in Fig. 2E). The equivalent residues in the putative E2 binding site of IFT172 are also depicted as sticks. (Right) Surface amino acid conservation map for the E2 binding site in IFT172 U-box domain, color-coded as indicated. **(F)** Western blot analysis of *in vitro* ubiquitination assay with HsIFT172C1 WT and the specified HsIFT172C1 U-box variants. Reactions were visualized by immunostaining with (top) anti-Ubiquitin antibody and (bottom) Coomassie staining.

Additional control experiments show that ubiquitin conjugates are only formed in the presence of all ubiquitination pathway components (E1, E2, ATP and ubiquitin) but not observed when WT UbcH5a is replaced by a catalytically dead mutant (Fig. 3D). Furthermore, the observed HsIFT172C1 ubiquitin conjugates disappear upon incubation with the Usp2 deubiquitinase enzyme and the HsIFT172C1 ubiquitin conjugate is resistant to reducing SDS-PAGE analysis (Fig. S3A and B). Combined, these observations indicate a canonical lysine-linked ubiquitin conjugation occurring on HsIFT172C1 in the presence of UbcH5a.

We were unable to test the U-box domain alone as the untagged HsIFT172C3 construct did not yield soluble protein expression. We did obtain protein expression for a GST-tagged HsIFT172C3 construct but several proteolytically cleaved fragments co-purified suggesting that the Ubox domain was partly degraded (Fig. S4B). Neither this GST-tagged HsIFT172C3 nor the HsIFT172C2 protein showed any ubiquitin conjugation (Fig. 3A). However, the longer CrIFT172_968-C_ and HsIFT172C1 did exhibit ubiquitin conjugation (Fig. 3A). Therefore, ubiquitination of HsIFT172 requires TPRs located N-terminally of the TPRs present in the crystallized construct shown in Fig. 2B, likely providing the ubiquitin accepting lysines.

We note that, unlike the canonical U-box/RING domain-containing proteins, IFT172 is not strongly poly-ubiquitinated *in vitro*. Moreover, Ubch5 proteins are highly reactive and often exhibit E3-independent ubiquitin transfer activity^63^ and it is thus possible that the observed IFT172 ubiquitination is a result of U-box-independent Ubch5 activity. As deletion of the U-box domain rendered IFT172 constructs insoluble in *E. coli*, we pursued mutagenesis studies on the U-box domain in context of the HsIFT172C1 construct. A common method to inactivate U-box/RING domains is to disrupt E2 binding^64^. Comparisons of the IFT172 sequence (Fig. 2F) and structure (Fig. 3E) with well-characterized U-box domains indicated two potential E2 binding residues in the IFT172 U-box domain. This includes a highly conserved Ile/Leu residue (I1688 in HsIFT172) in loop L1 of U-box/RING domains^51,64^ (Fig. 3E and highlighted with a red box in Fig. 2F) involved in E2 binding. Secondly, a conserved aromatic residue (F1715 in HsIFT172) is commonly found in helix α1 of U-box/RING domains^51,65^ (Fig. 3E and highlighted with a yellow box in Fig. 2F) where it contributes to E2 binding. Finally, a completely conserved proline residue is found in the loop L2 of U-box domains^51,53^ (P1725 in HsIFT172, see Fig. 2F and 3E), which in IFT172 caps the N-terminus of alpha-helix α2. Mutation of this conserved proline residue results in a complete loss of ubiquitination activity for four different U-box domain-containing proteins^53^. Interestingly, the IFT172 ciliopathy variant C1727R^36^ also maps to helix α2 in the U-box domain (Fig. 3E). Notably, I1688 and F1715 are located near an evolutionarily conserved surface patch on the putative E2 binding site of IFT172 (Fig. 3E). To test the importance of these residues in IFT172 ubiquitination, the residues Ile1688, Phe1715 and Pro1725 in IFT172 were mutated to alanine, whereas Cys1727 was mutated to arginine to mimic the patient variant. Soluble protein expression was obtained for each of the F1715A, P1725A, and C1727R HsIFT172C1 variants (but not for the I1688A variant) and the soluble proteins were purified (Fig. S3C-F). *In vitro* ubiquitination assays show that while mono-ubiquitination is not affected in IFT172 mutants, significant loss of the higher molecular weight bands for ubiquitinated IFT172 was observed for the F1715A and C1727R variants (Fig. 3F). IFT172 ubiquitin conjugation is thus affected by mutations of the putative E2 binding site of IFT172 U-box domain. We note that the strong mono-ubiquitination of IFT172 observed in Fig. 3F is likely U-box-independent ubiquitination by the E2 enzyme as this activity is not affected by IFT172 U-box mutations. In summary, we conclude that IFT172 exhibits ubiquitin conjugation in the presence of UbcH5a that is reduced in E2-binding and ciliopathy variants of IFT172. Therefore, while mutations in the IFT172 U-box domain affect the formation of higher molecular weight ubiquitin conjugates, the prominent mono-ubiquitination of IFT172 is likely attributable to the E3-independent activity of UbcH5a, as this event is not impacted by these U-box mutations, rather than indicating an intrinsic auto-ubiquitination capacity of IFT172 itself.

### IFT172 U-box domain pulls down ubiquitin *in vitro*

To further assess the biochemical properties of IFT172, we carried out interaction studies of HsIFT172C2 with stable mimics of the E2∼Ub (UbcH5a∼Ub) conjugate. To this end, the UbcH5a active site cysteine was mutated to a serine, which allowed for the production and purification of a stable oxyester-linked UbcH5a∼Ub conjugate^66^, which mimics the native thioester-linked E2-Ub conjugate (Fig. S4C). Interactions between IFT172 and the UbcH5a∼Ub conjugates were analyzed by affinity pull-downs using GST-tagged HsIFT172C2 (Fig. 4A) showing an interaction between IFT172C2 and the WT UbcH5a∼Ub conjugate. Additionally, we produced and tested two point-mutants of the UbcH5a∼Ub conjugate (UbcH5a F62A or A96D) known to disrupt interaction with U-box/RING domains^67–69^. However, these mutations did not prevent the interaction between UbcH5a∼Ub and HsIFT172C2 (Fig. 4A). Moreover, structural modeling of a hetero-dimeric complex between HsIFT172C3 and UbcH5a using AF-M did not indicate a direct interaction (Fig. 4B). *In silico* screening of complex formation between IFT172C3 and each of the 40 annotated Hs E2 enzymes also did not reveal confident complex formation with any of the E2 enzymes (data not shown). Combined, this indicates that although IFT172 associates with the E2-Ub conjugate in pull-downs, this association may not be mediated by direct interactions with the E2 enzyme (Fig 4A).

**Figure 4.**
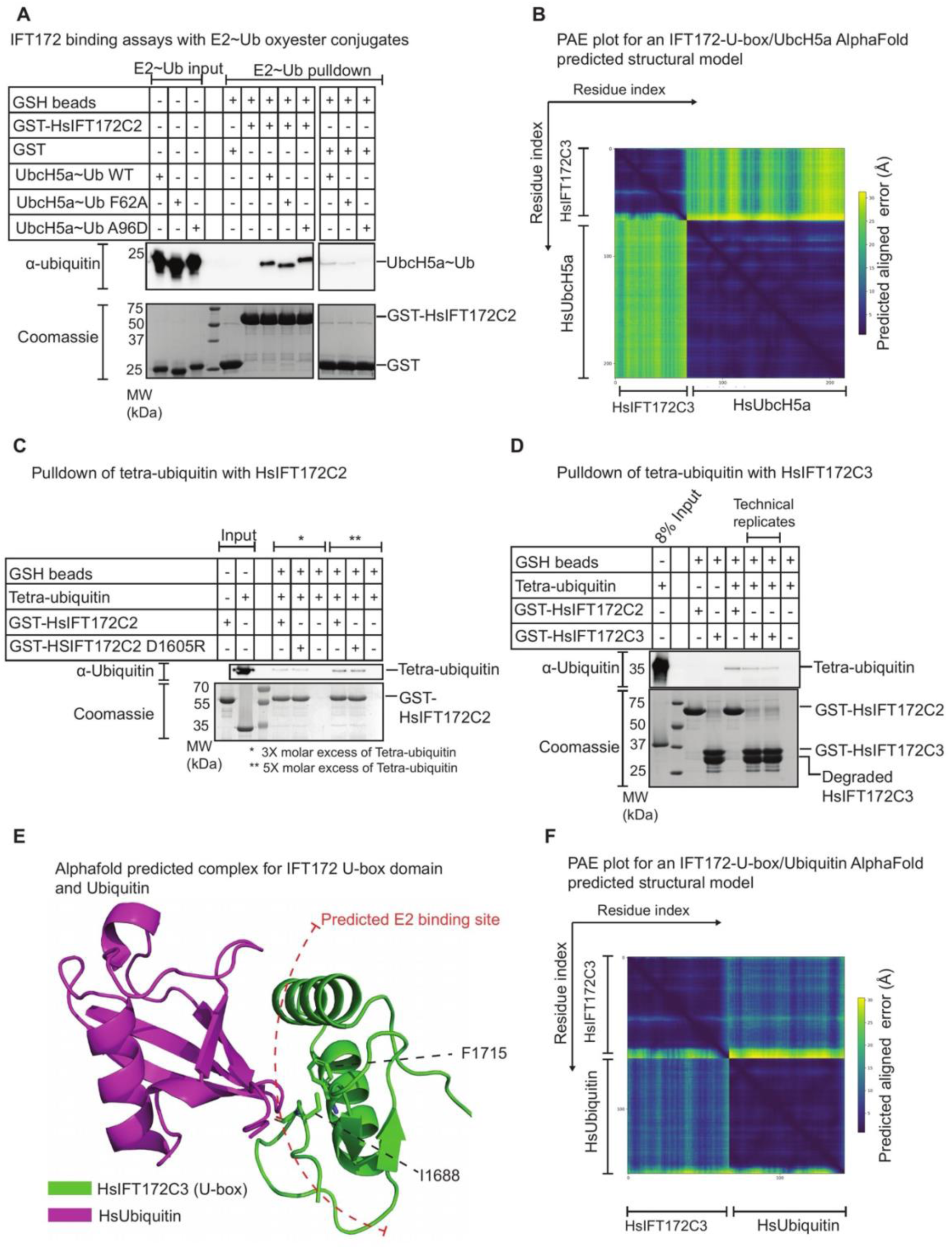
IFT172 U-box domain is a binding site for ubiquitin. **(A)** Pull-down of purified UbcH5a_C85S_∼Ub conjugates with GST-tagged HsIFT172C2 immobilized on GSH beads. Samples after elution were analyzed by western blotting with (top) anti-ubiquitin antibody and (bottom) Coomassie staining. Input shown is 2% for the western blot and 7.8% for the Coomassie staining. **(B)** PAE plot for an AlphaFold-generated structural model for a complex between HsIFT172C3 and HsUbcH5a. High error values are observed for interchain residue pairs, suggesting no high confidence prediction for an interaction between the two chains. **(C)** Pull-down of tetra-ubiquitin with GST-tagged HsIFT172C2 constructs immobilized on GSH beads. The D1605R mutation on the IFT-A binding site of HsIFT172C2 does not impact the binding of tetra-ubiquitin to HsIFT172C2. Reactions were visualized by immunostaining with (top) anti-ubiquitin antibody and (bottom) Coomassie staining. **(D)** Pull-own of tetra-ubiquitin with various GST-tagged HsIFT172 constructs immobilized on GSH beads. Both HsIFT172C2 and HsIFT172C3 pull-down tetra-ubiquitin at similar levels. The prominent lower molecular weight band in the HsIFT172C3 sample is a degradation/proteolytic cleavage product obtained upon HsIFT172C3 expression and purification from *E. coli*. The two lanes showing a pull-down of tetra-ubiquitin with HsIFT172C3 represent technical replicates. Reactions were visualized by immunostaining with (top) anti-Ubiquitin antibody and (bottom) Coomassie staining. **(E)** AlphaFold-predicted structural model for a complex between HsIFT172C3 and HsUbiquitin. The model suggests that ubiquitin binds to the predicted E2∼Ub binding site of HsIFT172C3. The predicted E2 binding residues of the HsIFT172 U-box domain are depicted in the model. **(F)** PAE plot for the AlphaFold structural model shown in panel E. Moderate PAE scores are observed for the residue pairs corresponding to the interaction interface (panel E) between the two chains.

Alternatively, the interaction between IFT172 and UbcH5a∼Ub could be mediated through ubiquitin. Indeed, several crystal structures have elucidated the RING-E2∼Ub interaction, demonstrating direct interactions between the RING domain and ubiquitin^55,56,58^. Given that E2 mutations do not abolish the interaction between UbcH5a∼Ub and HsIFT172C2, we reasoned that the interaction could be mediated by ubiquitin. N-terminally linked tetra-ubiquitin chains were pulled down with GST-tagged HsIFT172C2 indicating an IFT172-ubiquitin interaction (Fig. 4C). To test if the conserved IFT144 and IFT140 binding site on IFT172 is involved in ubiquitin binding, a pull-down was carried out with tetra-ubiquitin with the HsIFT172C2_D1605R mutant. Asp1605 corresponds to a previously reported human ciliopathy locus^37^ that maps onto the corresponding IFT144 and IFT140 binding site on IFT172. This HsIFT172C2_D1605R mutation does not appear to influence the binding to ubiquitin (Fig. 4C). Pull-down of the tetra-ubiquitin chain with GST-tagged HsIFT172C2 or HsIFT172C3 showed that both constructs retain tetra-ubiquitin-binding activity (Fig. 4D). The signal observed for HsIFT172C3 was somewhat reduced compared to HsIFT172C2, which likely reflects the partial proteolytic degradation of the GST-HsIFT172C3 construct during purification (Fig. S4B) rather than an intrinsic difference in ubiquitin-binding affinity. This suggests that the U-box domain likely contains the ubiquitin-binding site. Indeed, AF-M modeling of the IFT172 U-box domain with ubiquitin suggests the formation of a complex between the IFT172 U-box and ubiquitin (Fig. 4E-F). It was previously reported that E2 enzymes catalyze the E3-independent mono-ubiquitination of proteins containing a ubiquitin-binding domain^63^. It is thus possible that the mono-ubiquitination of HsIFT172C1 observed in the presence of UbcH5a (Fig. 3) could be a result of the ubiquitin-binding capability of HsIFT172C1.

### Loss of IFT172 U-box domain impairs protein stability and leads to ciliogenesis defects in RPE1 cells

To investigate the cilium-specific functions of the U-box domain of IFT172, we used CRISPR/Cas12a to engineer human RPE1 cell lines carrying a deletion of the IFT172 U-box domain. These cell lines were used to assess the impact of U-box domain loss on ciliogenesis, ciliary IFT172 localization, and TGFβ-1- and PDGF-DD-stimulated signaling responses (Fig. 5 and Fig. S5). A schematic overview and nomenclature for the four generated GFP-tagged RPE1 cell lines are shown in Fig. 5A, with a detailed description of the cell line generation strategy provided in the Materials and Methods section (Fig. S6A-B and Fig. S7A-B). To validate the engineered lines, we performed Sanger sequencing to confirm precise GFP knock-in and to verify the absence of undesired insertions or deletions (indels) (Fig. S6C and Fig. S7C). Expression of the tagged full-length and U-box-truncated IFT172 proteins was confirmed by immunoblotting with an anti-IFT172 antibody, which additionally revealed reduced steady-state levels of the IFT172ΔU-box protein compared to full-length IFT172, indicating that loss of the U-box domain compromises IFT172 protein stability (Fig. S8). Ciliogenesis in these cells was induced by serum withdrawal for 24h and cilia were visualized by staining for acetylated tubulin. In cells with a homozygous IFT172 U-box domain deletion cilia formation was severely reduced compared to cells expressing GFP-tagged full-length IFT172 (Fig. 5B). Moreover, cilia that are retained in the IFT172ΔU-box (homozygous) cells are significantly shorter, with a median cilia length of ∼1.3 µm in IFT172ΔU-box (homozygous) cells, compared to ∼3.0 µm in IFT172-FL (homozygous) cells (Fig. 5B). Meanwhile, in cells with a heterozygous IFT172 U-box domain deletion no significant effect on ciliation or ciliary length was observed (Fig. 5B). To further investigate the observed ciliogenesis phenotype, the localization of the eGFP-tagged IFT172 proteins were analyzed by immunofluorescence microscopy along with quantification of fluorescence signal along the ciliary axis. Both the full-length IFT172 protein as well as the IFT172ΔU-box truncation were recruited to the cilium (Fig. 5C and S5C-F). The full-length IFT172 protein is present along the length of the axoneme, with notable accumulations at the ciliary base and tip. In the cells with a homozygous IFT172 U-box deletion, eGFP-tagged IFT172ΔU-box protein is accumulated along the short axoneme (Fig. S5E), but in heterozygous cells that form normal length cilia, the distribution of the eGFP-tagged IFT172ΔU-box protein is similar to the full-length IFT172 protein (Fig. 5C, Fig. S5D and F). This consistent localization pattern suggests that the U-box domain is not essential for IFT172 trafficking to or within the cilium, despite its impact on overall protein levels and ciliogenesis.

**Figure 5.**
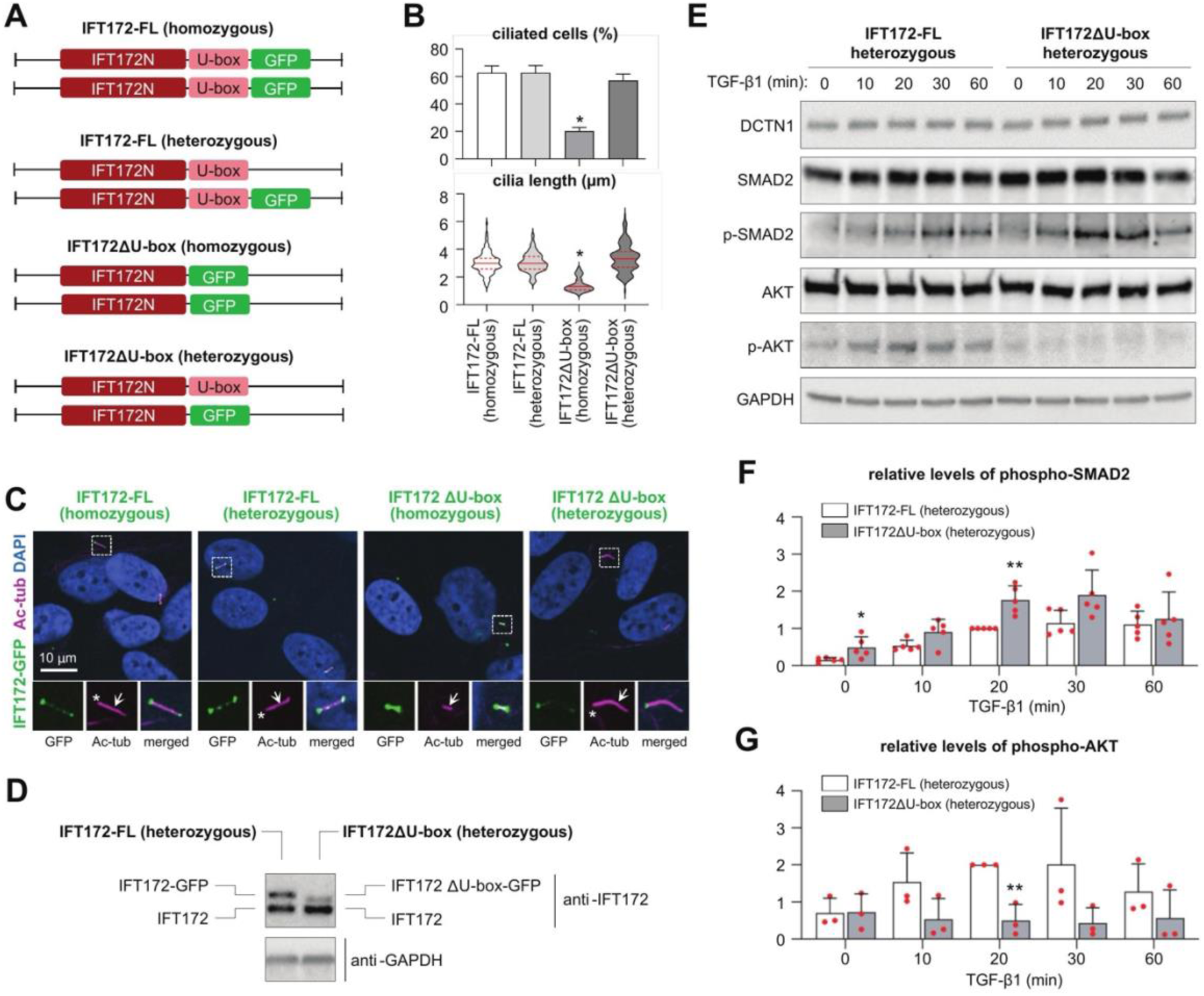
Truncation of the IFT172 U-box domain impairs ciliogenesis and leads to altered TGF-β signaling response in RPE1 cells. **(A)** Schematic representation and nomenclature of RPE1 cell lines generated by CRISPR/Cas12a-mediated exon targeting of the IFT172 gene. **(B)** Quantification of ciliogenesis (percentage of ciliated cells, upper panel) and ciliary length (lower panel) in all RPE1cell lines. Error bars in the upper panel represent standard error of the mean (*: p<0.05). **(C)** IFM analysis of IFT172-GFP (*green*) localization to primary cilia (acetylated tubulin (Ac- tub., *purple*) in RPE1 cell lines. Nuclei are stained with DAPI (*blue*). The top row displays whole-cell views, while the bottom row panels show zoomed in insets of the cilium (arrows). Asterisks indicate ciliary base region. **(D)** WB analysis of IFT172 expression in IFT172-FL (heterozygous) and IFT172ΔU-box (heterozygous) RPE1 cell lines. **(E)** WB analysis of phosphorylation levels of SMAD2 (p-SMAD2) and AKT (p-AKT; p-AKT^T308^) in IFT172-FL (heterozygous) and IFT172ΔU-box (heterozygous) RPE1 cell lines treated with 2 ng/mL TGF-β1 ligand for the indicated time points. **(F,G)** Quantification of p-SMAD2 (F) and p-AKT (G) levels from panel E, normalized to DCTN1 and GAPDH. Error bars represent standard error of the mean (*: p<0.05; **:p<0.01).

Examination of protein expression levels using immunostaining revealed a striking reduction in the levels of the U-box deleted IFT172 protein (Fig. 5D). The specific IFT172 antibody used here was generated against an HsIFT172 epitope (amino acids 1353-1652) that lies outside the U-box domain. The reduced detection of IFT172 is thus most likely due to reduced protein expression or stability levels upon U-box deletion. As IFT172 is known to be essential for ciliogenesis^41,70^, the severe reduction in expression levels of both IFT172 alleles causes ciliogenesis defects in the IFT172ΔU-box (homozygous) cells. Meanwhile, in the IFT172ΔU-box (heterozygous) cells, the observed upregulation in the WT IFT172 allele compensates for the reduced expression levels of the U-box truncated IFT172 allele (Fig. 5D), which likely provides sufficient IFT172 for ciliogenesis at a similar level as the parental RPE1 cells (Fig. 5B). However, the IFT172ΔU-box protein contains both IFT-A and IFT-B binding sites required for incorporation into IFT trains and the IFT172ΔU-box (heterozygous) cells will thus have a reduction in the concentration of IFT172 with a U-box domain as compared to parental cells.

### IFT172 U-box domain influences TGF-Β signaling

Given that the U-box deleted IFT172 still localizes to the cilium (Fig. 5C), we wanted to investigate if ciliary signaling events are impacted by the loss of the IFT172 U-box domain. The severe ciliogenesis defects exhibited by the IFT172ΔU-box (homozygous) cells (Fig. 5B) will result in U-box-independent secondary effects that may affect signaling responses. Therefore, we focused our attention on the IFT172ΔU-box (heterozygous) cells that possess normal ciliation and initially investigated TGF-Β signaling in these mutant cells. To this end, previous studies showed that the primary cilium regulates both canonical and non-canonical branches of TGF-β signaling^71,72^ the former operating through activation of R-SMAD transcription factors at the ciliary pocket, where ciliary receptors are internalized for phosphorylation of SMAD2/3^71^. Compared to control cells, IFT172ΔU-box (heterozygous) cells showed an increased level of SMAD2 activation upon TGF-β1 stimulation, and the level of SMAD2 phosphorylation also appeared slightly elevated in unstimulated heterozygous IFT172ΔU-box cells (Fig. 5E-F). In contrast, TGF-β1 mediated activation of AKT, which operate in the non-canonical branch of TGF-β signaling, was markedly reduced in heterozygous IFT172ΔU-box cells (Fig. 5E and 5G). We further evaluated AKT activation in response to PDGF-DD stimulation, which activates the homodimer of PDGFRB outside and independent of the primary cilium^73^. In this case, PDGF-DD induced activation of AKT was unaffected in the heterozygous IFT172ΔU-box cells (Figs. S5A-B). Thus, perturbation to the IFT172 U-box domain results in differential effects on TGF-β mediated signaling pathways, for which the U-box domain may play a regulatory role in the mechanisms by which the cilium balances the output of canonical versus non-canonical TGF-β pathways.

## Discussion

### IFT172 bridges IFT-A and IFT-B complexes in IFT trains

This study provides new insights into the structure and function of IFT172, a key component of the intraflagellar transport machinery. The N-terminal region of IFT172 is known to form inter-IFT subunit interactions with the IFT-B subunits IFT57/IFT80^34,35,74^, while much less was known about the C-terminal part of IFT172. Our comprehensive structural and biochemical analyses reveal two functionally relevant motifs at the C-terminus of IFT172: a TPR motif involved in IFT-A association and a U-box-like domain likely involved in ciliary ubiquitination events. These discoveries not only enhance our understanding of the role of IFT172 in IFT but also suggest new mechanisms for the regulation of ciliary function.

Early studies on IFT172 hypothesized a function in anterograde to retrograde transition of IFT trains at the ciliary tip^40^. This hypothesis was based on the observation that the *Chlamydomonas reinhardtii fla11* strain with a C-terminal missense mutation in IFT172 leads to an accumulation of IFT proteins at the ciliary tip, a phenotype characteristic of retrograde IFT defects^40^. Our findings now provide a molecular basis for the retrograde IFT phenotype observed in the *fla11* strain. We show that the FLA11 mutation lies in the TPR motif of IFT172 directly associating with IFT subunits IFT144 and IFT140 (Fig. 1 and S1). This suggests that impairment of the association of IFT172 with IFT-A likely underlies the faulty switch to retrograde IFT. Our results align with recent cryo-ET structures of anterograde and retrograde IFT trains, which show that the IFT172 C-terminus interacts with IFT144 in anterograde trains and with IFT140 in retrograde trains^46,47^. While L1615P is predicted to affect both interactions, the predominantly retrograde IFT phenotype of fla11 can be rationalized by the distinct roles of the IFT172 C-terminus in anterograde versus retrograde trains. In anterograde trains, the IFT172 C-terminus functions as a flexible tether in stoichiometric excess, where additional lateral interactions between IFT-B subunits provide redundancy that likely compensates for reduced affinity caused by L1615P46. In contrast, the retrograde train requires the IFT172 C-terminus to adopt a rigid conformation integral to the IFT-A dimeric interface, with no compensatory lateral interactions^47^. The helix-breaking L1615P mutation would thus selectively disrupt retrograde train assembly.

Interestingly, the human ciliopathy variant D1605E^37^ also localizes to the corresponding IFT-A interaction site in the *Hs*IFT172, suggesting that impaired IFT-A association contributes to disease manifestation. The additional methylene group in the glutamate side chain may introduce steric clashes at the tightly packed IFT172-IFT-A interface, providing a rationale for the pathogenicity of this conservative substitution. Collectively, these findings establish IFT172 as a key IFT subunit that bridges IFT-B and IFT-A complexes in both anterograde and retrograde IFT trains, providing a molecular basis for its involvement in ciliopathies.

We note that IFT-A subunits IFT121, IFT122, and IFT144 share a similar domain organization with IFT172, with TPR repeats terminating in small C-terminal Zn-finger-like domains^95^. Structural superposition reveals a shared topology, raising the possibility that these domains share a common evolutionary origin. While this could suggest conserved ubiquitin-related functions across these subunits, the more parsimonious explanation is that these C-terminal domains primarily serve structural roles in protein stabilization or IFT subunit interactions. Indeed, we cannot exclude that the U-box-like domain of IFT172 similarly contributes to protein stability, as evidenced by the reduced IFT172 levels upon U-box deletion (Fig. 5D). Whether the U-box-like domain of IFT172 has acquired an additional ubiquitin-related function beyond its structural role remains an open question requiring further investigation.

### Stoichiometry and potential ubiquitin-related functions of IFT172 in IFT trains

Recent cryo-EM reconstructions of IFT trains have revealed a stoichiometry of 2:1 for IFT-B to IFT-A complexes^46,47^. This stoichiometry implies that only half of the IFT172 C-termini are engaged with IFT-A, while the remainder are potentially free to interact with other ciliary components. Our discovery of a U-box-like domain in IFT172 (Fig. 2), and the identification of a UBX domain protein as an interaction partner, suggests potential ubiquitin-related functions for IFT172 within the cilium. These functions could extend beyond proteasomal degradation to include roles in ubiquitin-mediated protein trafficking or regulation. This hypothesis is supported by our observation of weak *in vitro* ubiquitin-conjugation of IFT172 (Fig. 3F). Further demonstration of ubiquitination activity of IFT172 directly in cells by identifying *bona fide* ubiquitination substrates is required to assign the ubiquitin ligase function to IFT172.

The IFT172 U-box domain appears to be in an auto-inhibited state in our crystal structure of HsIFT172C2 (Fig. 2E), potentially explaining the absence of a robust auto-ubiquitination activity *in-vitro*. This structural inhibition is reminiscent of the RING ubiquitin ligase CBL^59^, where phosphorylation and substrate binding trigger a conformational change that activates ligase activity^59,75^. Intriguingly, the phosphosite database^76^ lists four residues (T1533, S1549, T1689, Y1691) at the U-box/TPR interface as phosphorylation sites (Fig. S2D). Phosphorylation of these residues could potentially alleviate the auto-inhibited state, suggesting a possible regulatory mechanism. Furthermore, a 30-residue linker connects the U-box domain to the last TPR of IFT172, likely providing significant conformational flexibility (Fig. 2A-B). This flexibility may be functionally crucial for the U-box domain, allowing it to adopt different conformations as needed for its various roles. However, we note that the proposed autoinhibition model and its potential regulation by phosphorylation remain hypothetical and require future experimental validation.

Our biochemical characterization has further revealed ubiquitin-binding properties of IFT172, mapped to the U-box-like domain (Fig. 4). While this property is atypical for U-box domains, it bears resemblance to structurally related zinc finger domains, such as the UBZ domain with a β-β-α motif^77^. These domains function as ubiquitin-binding domains (UBDs) regulating various cellular processes^77^. This finding is particularly intriguing as RING domains, which are structurally similar to U-box domains, are known to mediate contacts with ubiquitin for priming the E2-Ub conjugate for ubiquitin transfer^55,56^.

The ubiquitin-binding capability of IFT172 could facilitate the trafficking or turnover of ubiquitinated proteins in the cilium, complementing known ubiquitin-dependent processes. For instance, signal-dependent ciliary exit of ubiquitinated GPCRs and the Hedgehog pathway component Smoothened^25,29^ is mediated by UBDs in the adaptor protein TOM1L2^26,31^. However, UBDs associated with several other cilium-associated signaling pathways and processes are yet to be identified. While IFT139 has been implicated in the ciliary turnover of ubiquitinated tubulin during ciliary disassembly^28^, unlike TOM1L2, ubiquitin-binding sites have not been validated in IFT139 or any other BBSome or IFT subunits. This positions the IFT172 U-box domain as a potential UBD for the trafficking or turnover of ubiquitinated proteins in the cilium.

### Implications of the IFT172 U-box domain in ciliary signaling

Several ciliopathy variants map to the C-terminal part of IFT172^36–38^ and are located in the vicinity of the U-box domain (Fig. 2B). These mutations could disrupt ubiquitin-related functions of the U-box domain or impair protein stability. To investigate the physiological relevance of the IFT172 U-box domain, we analyzed engineered RPE1 cells with U-box deletions. While homozygous deletion of the U-box domain significantly impaired total IFT172 protein levels and affected ciliogenesis, a heterozygous deletion maintained near-wild-type levels of total IFT172 protein and cilium formation (Fig. 5B-D). In the heterozygous U-box deleted cells, we observed altered SMAD2 phosphorylation levels (Fig. 5E-F) and impaired AKT activation (Fig. 5E and G) in response to TGF-β1 stimulation. These findings suggest that the U-box domain is crucial for protein stability and its deletion differentially affects the SMAD and non-SMAD TGF-β activation pathways. The final phenotypic outcome of TGF-β signaling depends on the balance and fine-tuning of both SMAD and non-SMAD activation pathways, as these pathways engage in extensive cross-talk and can mutually regulate each other^78–80^. Our results suggest that IFT172, through its U-box domain, may play a crucial role in maintaining this balance. However, we note that our current experiments cannot distinguish whether these signaling effects result specifically from loss of ubiquitin-related functions of the U-box domain or from the reduced levels of functional IFT172 protein in the heterozygous U-box deleted cells. Targeted point mutations that selectively disrupt ubiquitin binding without affecting protein stability would be required to resolve this question. We further note that our conclusions are based on a single validated clone per genotype, and confirmation in independent clones will be important to fully exclude clonal effects.

While ciliary regulation of SMAD2/3 phosphorylation via TGFβ receptors is well established, the connection between primary cilia and AKT signaling is also increasingly supported. AKT has been shown to regulate ciliogenesis initiation through a Rab11-effector switch mechanism, where disruption of AKT signaling promotes the transition from Rab11-WDR44 to the ciliogenic Rab11-FIP3-Rabin8 complex^96^. Furthermore, a cilia-dependent reciprocal activation between AKT1 and SMAD2/3 has been demonstrated in trabecular meshwork cells, where primary cilia act as mechanosensors regulating both pathways^97^. Our observation that heterozygous deletion of the IFT172 U-box domain differentially affects SMAD2 phosphorylation and AKT activation is consistent with the emerging role of primary cilia in coordinating canonical and non-canonical TGFβ signaling, although the precise mechanism by which the U-box domain influences these pathways requires further investigation.

In conclusion, our findings reveal that IFT172, beyond its structural role in IFT trains, has an unexpected function in signal regulation. The flexible nature of half of the IFT172 C-termini in IFT trains, combined with its ubiquitin-related activities, positions IFT172 as a potential regulator of ubiquitin-mediated processes within the cilium. This dual functionality - providing both structural support and signaling regulation - may explain why IFT172 mutations lead to such diverse ciliopathy phenotypes. These results expand our understanding of how IFT proteins contribute to ciliary signaling beyond their established roles in protein transport.

## Materials and Methods

### Cloning and expression of proteins in *E.coli*

DNA sequences encoding the respective truncations of HsIFT172 and CrIFT172 as well as full length HsUbiquitin and HsUbcH5a were Polymerase Chain Reaction (PCR) amplified. Gibson assembly^81^ was used to insert the genes into pEL_A or pEL_K vectors with either N-terminal 6XHis-TEV or 6XHis-GST-TEV tags. Mutations were introduced by PCR of the corresponding plasmids. Plasmids were transformed into *E. coli* BL21 (DE3) and the cells were grown at 37°C in TB medium supplemented with appropriate antibiotics to an OD_600_ of 1. After cooling down the culture to 18°C, protein expression was induced by addition of 0.5 mM Isopropyl β- d-1-thiogalactopyranoside (IPTG). Cells were harvested after overnight protein induction at 18°C.

### Protein purification

Cell pellets were resuspended in four times the pellet volumes of lysis buffer (50 mM Tris pH 7.5, 300 mM NaCl, 10% glycerol, 5 mM β-mercaptoethanol) supplemented with 1 mM phenylmethanesulfonyl fluoride (PMSF) and SM DNAse. The cells were lysed by sonication and clarified by ultracentrifugation at 74000 relative centrifugal force (RCF) for 30 minutes. The cleared lysate was loaded onto a Ni^2+^-NTA column (5 ml, Roche) prewashed with lysis buffer. After loading the lysate, the column was further washed with 8 Column volumes (CV) of lysis buffer, 8 CV QB buffer (20 mM Tris pH 7.5, 1 M NaCl, 10% glycerol, 5 mM β-mercaptoethanol) and 8CV of QA buffer (20 mM Tris pH 7.5, 50 mM NaCl, 10% glycerol, 5 mM β-mercaptoethanol) supplemented with 20 mM Imidazole. The proteins were eluted from the Ni^2+^-NTA column by passing QA buffer containing 250 mM imidazole. The eluted proteins were loaded onto a HiTrap Q HP 5 ml anion exchange column (GE Healthcare) pre-equilibrated with QA buffer. Elution from the Q column was performed by a 0-100% gradient from QA to QB buffer. The elutions containing the proteins were loaded onto a HiLoad 16/600 Superdex 75 or HiLoad 16/600 Superdex 200 column (GE Healthcare) pre equilibrated with SEC buffer (10 mM HEPES pH 7.5, 150 mM NaCl, 1 mM Dithiothreitol (DTT)).

For the purification of His-GST-TEV-HsIFT172_1681-C_, His-MmUbe1, His-TEV-HsUbcH5a, His-TEV-(Tetra-ubiquitin) and His-Strep-TEV-HsUbiquitin the cell lysates over expressing the proteins were prepared as above. After loading the clarified lysate onto the Ni^2+^-NTA column, the column was washed with 8CV of lysis buffer, 8 CV of QB buffer and 8CV of QA2 buffer (20mM Tris pH 7.5, 100mM NaCl, 10% glycerol, 5 mM β-mercaptoethanol) supplemented with 20mM Imidazole. Proteins were eluted by passing 20mL QA2 buffer containing 250mM Imidazole followed by 20mL of QA2 buffer containing 500mM Imidazole. The purest elutions were dialyzed overnight in SEC buffer and afterwards loaded on a HiLoad 16/600 Superdex 75 or HiLoad 16/600 Superdex 200 column (GE Healthcare) pre-equilibrated with SEC buffer. Selenomethionine substituted HsIFT172C2 protein was expressed in BL21 Star cells and grown in M9 medium with addition of amino acids. The protein was purified as described above.

### Culturing and flagella isolation of Cr

CrCC-1690 strain was obtained from the Chlamydomonas Resource Center (https://www.chlamycollection.org). The cells were maintained on solid Agar plates consisting of standard tris acetate phosphate (TAP) media in a sterile environment at room temperature with a table lamp as a continuous light source. Cells for flagella extraction were grown in two flasks each with 4L of (TAP) media for 3 days at room temperature with a table lamp as continuous light source. Cells were grown to an OD_700_ of 0.4. The cultures were harvested and dibucaine induced flagella abscission and subsequent flagella isolation was carried out as reported in Craige et al., 2013^82^. The isolated flagella were solubilized overnight at 4°C in solubilization buffer (10mM HEPES pH 7.5, 5 mM MgSO_4_, 4% Sucrose, 25mM KCl, 10mM β-mercaptoethanol, 0.3% IGEPAL 630) supplemented with protease inhibitor (Roche #05892791001). Afterwards, the axonemal fraction of the flagella lysate was pelleted by centrifugation of the flagella lysate at 45,000 RCF for 30min at 4°C. The supernatant containing the IFT proteins and motor proteins were used for subsequent pull-downs with CrIFT172_968-C_.

### Affinity pull-down

For the pull-down with *Chlamydomonas reinhardtii* flagella extracts, 30μM of His-crIFT172_968-C_ or 30μM His-TEV protease were immobilized on 50μL of TALON beads (GE healthcare #28-9574-99) by incubation at 4°C for 1 hour. Afterwards the beads were washed 3 times with HMSK buffer (10mM HEPES pH 7.4, 5mM MgSO_4_, 4%(w/v) sucrose and 25mM KCl) to remove unbound proteins. 1mg *Chlamydomonas reinhardtii* flagella extract diluted in 300μL HMSK buffer was incubated with the beads at 4°C for 2 hours. After incubation, beads were washed three times with HMSK buffer containing 30mM Imidazole and bound proteins were eluted in HMSK buffer containing 300mM Imidazole.

For the GST pull-down 10μM of the GST tagged protein or GST tag were immobilized on 10 μL GSH beads (Cytiva #17-5279-01) by incubation for 1 hour at 4°C with constant mixing. Beads were washed once with pull-down buffer (50mM Tris pH 7.5, 100mM NaCl and 1mM DTT) to remove unbound proteins. Prey proteins were diluted in 100μL pull-down buffer and incubated with the beads at 4°C for 2 hours. The beads were washed three times with pull-down buffer and bound proteins were eluted in elution buffer (10mM HEPES pH7.5, 100mM NaCl, 30mM Reduced glutathione and 1mM DTT). The specific amount of prey proteins supplied were 20μM UbcH5a∼Ub in the pull-down shown in Fig. 4A, 25μM tetra-ubiquitin in the pull-down shown in Fig. 4D. For the pull-down shown in Fig 4C, 5μM of immobilized GST-HsIFT172C2 was incubated with 15μM or 25μM tetra-ubiquitin as specified. Pull-down elutions were visualized by Coomassie staining and western blotting with anti-ubiquitin (Merck, #05-944, 1:5000) antibody after separation on an SDS-PAGE gel.

### Ubiquitination assays

Ubiquitination assays with purified UbcH5a as E2 were performed in a 50 μL reaction volume. The reaction contained 0.1 μM E1 (His-MmUbe1), 2.5 μM E2 (His-TEV-UbcH5a), 10 μM ubiquitin (R&D Systems #U-100H-10M) and 2 μM of the specified His-TEV-HsIFT172C constructs. Reactions were initiated by addition of 5 mM ATP and incubated at 37°C for 1.5 hours in ubiquitination buffer (50 mM Tris pH 7.5, 50 mM NaCl, 5 mM MgCl_2_ and 0.5 mM DTT). The reactions were stopped by addition of equal volumes of 2X SDS-PAGE loading buffer (100 mM Tris pH 6.8, 10% v/v β-mercaptoethanol,4% v/v SDS, 0.2% v/v bromophenol blue, 20% v/v glycerol). Reactions were later visualized by Coomassie staining and western blotting with anti-ubiquitin (Merck,#05-944, 1:5000) antibody or anti-His (GeneScript, #A00186, 1:5000) antibody after separation on an 8% SDS-PAGE gel.

For the E2 enzyme screen shown in Fig. 3B, 20 μL reactions were set up containing 0.25 μM E1 (His-MmUbe1), 2.5 μM of the specified E2 enzyme (Abcam #ab139472), 5 μM ubiquitin (R&D Systems #U-100H-10M) and 2 μM His-TEV-HsIFT172C1. The reactions were initiated by addition of 5 mM ATP and carried out in ubiquitination buffer (Abcam #ab139472) supplemented with 5 mM MgCl_2_. The reactions were incubated at 37°C for 1.5 hours. The reaction was stopped by addition of equal volumes of 2X non-reducing SDS-PAGE loading buffer (Abcam #ab139472). Reactions were analyzed by Coomassie staining and western blotting with anti-ubiquitin antibody (Merck, #05-944, 1:5000) after separation on a 4-15% gradient SDS-PAGE gel.

The protocol for generating the UbcH5aC85S∼Ub conjugate (Fig. S4C), was adapted from Middleton et al., 2014^66^. Briefly, a 5 mL *in-vitro* charging reaction was set up containing MmUbe1, UbcH5a_C85S_, ubiquitin (recombinantly purified with an N-terminal 6XHis-Strep-TEV tag) and ATP. The charging reaction was incubated overnight at 30°C to achieve maximal UbcH5aC85S∼Ub formation, followed by separation on a HiLoad Superdex 75 SEC column.

### Cell culture

Adherent hTERT-immortalized Retinal Pigment Epithelial 1(RPE1) cells were grown in Dulbecco’s Modified Eagle Medium F12 (DMEM/F-12, GlutaMAX Supplement (Gibco #31331-093)) containing 1% penicillin-streptomycin (Sigma-Aldrich #P0781) and 10% fetal bovine serum (FBS Gibco #10438-026) at 37°C, 5% CO_2_ and 95% humidity. Cells were passaged twice a week. Cells were serum starved for 48 hours prior to ligand stimulation by replacing the culture media with DMEM F12 containing 1% Penicillin-Streptomycin.

### Ligand stimulation assays

Serum starved RPE1 cells and fibroblasts were stimulated by addition of respective serum starvation media containing 2ng/mL TGF-β1 ligand (R&D Systems #240B) for the specified time points. The stimulation was quenched by washing the cells with ice cold Phosphate-Buffered Saline (PBS) (137mM NaCl, 2.7mM KCl, 10mM Na_2_HPO_4_ and 1.8mM KH_2_PO_4_) followed by addition of lysis buffer. M-Per Lysis buffer (ThermoScientific #78501) supplemented with protease inhibitor (Merck #05056489001) and anti-phosphatase (ThermoScientific #1862495) was used for quenching and cell lysis. Lysates were centrifuged at 20000 RCF for 20 min at 4°C and the supernatant was stored at −20°C. Samples were later analyzed by western blots with antibodies as indicated. Antibodies used in western blots are listed in Table 2.

**Table 2:**
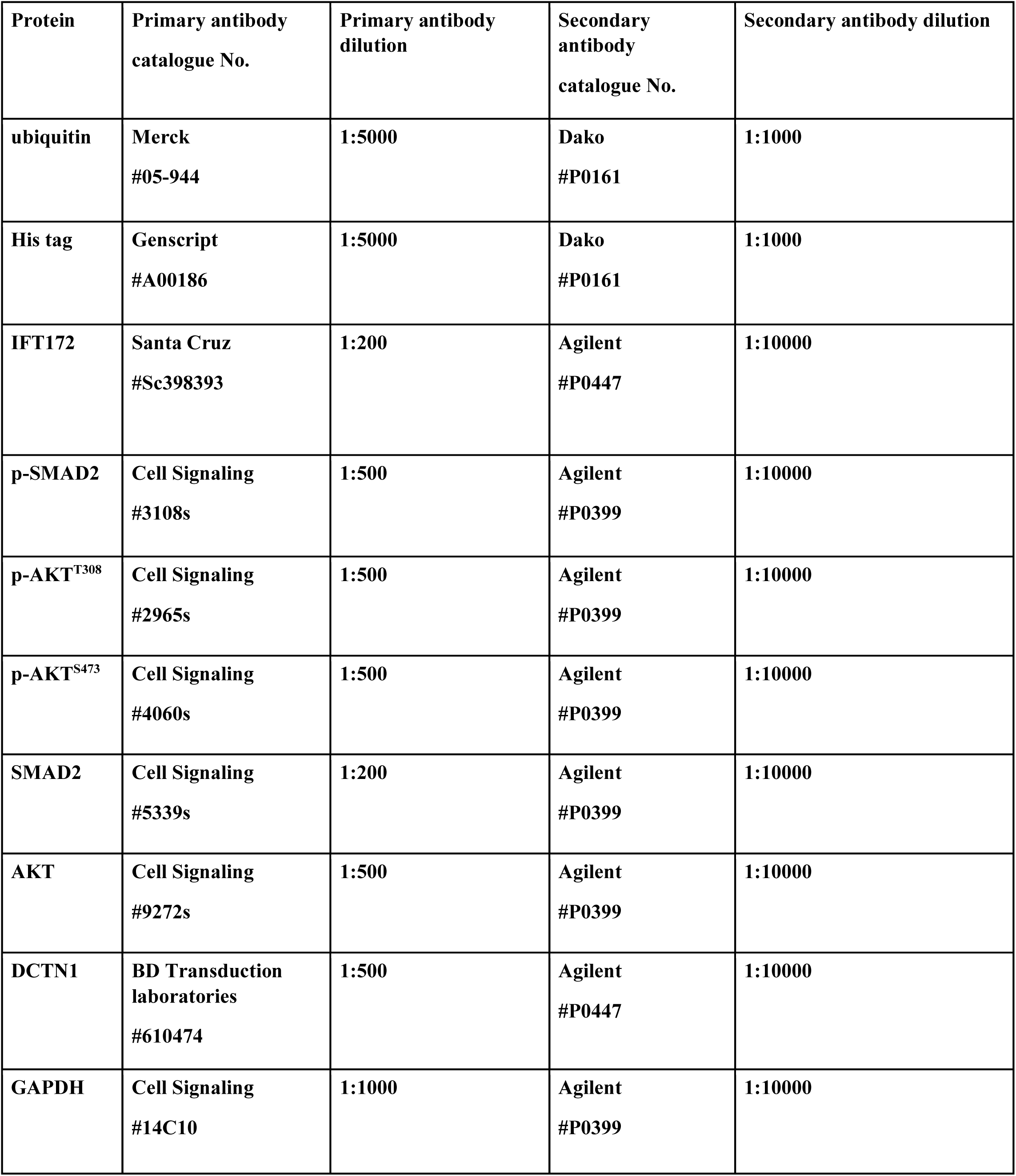
List of primary and secondary antibodies used in western blots.

| **Protein** | **Primary antibody catalogue No.** | **Primary antibody dilution** | **Secondary antibody catalogue No.** | **Secondary antibody dilution** |
| --- | --- | --- | --- | --- |
| **ubiquitin** | **Merck**<br>**#05-944** | **1:5000** | **Dako**<br>**#P0161** | **1:1000** |
| **His tag** | **Genscript**<br>**#A00186** | **1:5000** | **Dako**<br>**#P0161** | **1:1000** |
| **IFT172** | **Santa Cruz**<br>**#Sc398393** | **1:200** | **Agilent**<br>**#P0447** | **1:10000** |
| **p-SMAD2** | **Cell Signaling**<br>**#3108s** | **1:500** | **Agilent**<br>**#P0399** | **1:10000** |
| **p-AKT<sup>T308</sup>** | **Cell Signaling**<br>**#2965s** | **1:500** | **Agilent**<br>**#P0399** | **1:10000** |
| **p-AKT<sup>S473</sup>** | **Cell Signaling**<br>**#4060s** | **1:500** | **Agilent**<br>**#P0399** | **1:10000** |
| **SMAD2** | **Cell Signaling**<br>**#5339s** | **1:200** | **Agilent**<br>**#P0399** | **1:10000** |
| **AKT** | **Cell Signaling**<br>**#9272s** | **1:500** | **Agilent**<br>**#P0399** | **1:10000** |
| **DCTN1** | **BD Transduction laboratories**<br>**#610474** | **1:500** | **Agilent**<br>**#P0447** | **1:10000** |
| **GAPDH** | **Cell Signaling**<br>**#14C10** | **1:1000** | **Agilent**<br>**#P0399** | **1:10000** |

### Crystallization, X-ray diffraction data processing and structure determination

HsIFT172C2 was crystallized by vapor diffusion by mixing 200 nL of purified protein at a concentration of 2.4 mg/ml mixed with an equal volume of precipitant solution containing 0.2 M Sodium phosphate dibasic dihydrate, 20% w/v Polyethylene glycol 3350, pH 9.1. Crystals were transferred to the cryo-protectant containing the precipitant solution supplemented with 10% glycerol before freezing. X-ray diffraction data were collected at the Swiss Light Source (SLS; Villigen, Switzerland) at the PXII beamline on a Pilatus 6M detector and indexed/integrated with the XDS package^83^ before scaling with Aimless as part of the CCP4 package^84^. Molecular replacement using the AlphaFold generated model for HsIFT172C2 was carried out in the program Phaser^85^ as available in the software packages PHENIX^86^. Molecular replacement identified two molecules of HsIFT172_1470-C_ in the asymmetric unit. Single anomalous dispersion diffraction data collected at the Selenium peak wavelength on Selenium methionine substituted protein crystals were combined with the molecular replacement solution in Phaser to produce phase information to calculate the map shown in Fig. S2B. Given this map, the AutoBuild function in PHENIX was utilized for model building yielding initial R_work_/R_free_ values of 0.30/0.35. This was followed by iterative cycles of manual model building in Coot^93^ and refinement in PHENIX using torsion angle non crystallographic symmetry and secondary structure restraints as well as translation libration screw (TLS) refinement, to yield a final model with an R_work_/R_free_ of 0.195/0.240 (see Table 1).

### Mass Spectrometry based fragment identification for CrIFT172_968-C_ interactors

Samples were prepared using the SP3 protocol^87^ with reduction, alkylation, and trypsin digestion. Peptides were labeled with TMT6plex (ThermoFisher) and fractionated by high pH reverse phase chromatography. LC-MS/MS analysis was performed on an UltiMate 3000 RSLC nano LC system coupled to an Orbitrap Fusion Lumos Tribrid Mass Spectrometer. Full scan MS1 (375-1500 m/z) was acquired at 60,000 resolution, followed by data-dependent MS2 scans at 15,000 resolution. Data were processed using IsobarQuant and Mascot (v2.2.07) against the Chlamydomonas reinhardtii proteome (UP000006906). Fixed modifications were Carbamidomethyl (C) and TMT10 (K); variable modifications were Acetyl (Protein N-term), Oxidation (M), and TMT10 (N-term). Mass tolerances were 10 ppm for MS1 and 0.02 Da for MS2. Quantification required at least two unique peptides per protein. Data analysis was performed in R, using limma for batch correction and vsn for normalization. Differential expression was assessed using limma, with hits defined as having FDR < 5% and fold-change ≥ 100%, and candidates as FDR < 20% and fold-change ≥ 50%.

### CRISPR-mediated endogenous tagging of IFT172 in RPE1 cells

The CRISPR/Cas12a-assisted PCR tagging approach was used to endogenously tag IFT172 with eGFP in RPE1 cell line as previously described^88^ Briefly, HDR repair templates were produced by PCR with target-specific primers containing the homology arms and the plasmid pMaCTag-P05 (Addgene plasmid 120016)^89^ In addition to the homology arms and the eGFP sequence, the HDR repair template also encodes a puromycin cassette for selection and an expression cassette for a Cas12a crRNA. For tagging full-length IFT172 (IFT172-FL) at its C-terminus, the Cas12a crRNA (5’-CCTTTCAGTAGTTGGTAGAG-3’) targeted the IFT172 locus at the stop codon, and the homology arms were designed to insert the eGFP sequence before the stop codon. To generate the deletion of the IFT172 U-box domain (IFT172 ΔU-box), the homology arms were designed to insert the eGFP sequence after amino acid 1688, and the target sequence of the corresponding Cas12a crRNA (5’-TTACAGGTATAGAAGCCTAC-3’) spanned the intended integration site. Doxycycline-inducible RPE1 cells expressing the CRISPR nuclease enAsCas12a^90^ were electroporated with HDR repair template using the Neon Transfection System (ThermoFisher Scientific). After the transfection, the cells were treated for three days with the DNA-PK inhibitor M3814 to increase the knock-in efficiency^91^ and subsequently selected with puromycin for 2 weeks, to isolate single cell clones. The clonal cell lines were screened for successful tagging by live cell imaging and PCR. In this study four cell lines were used (Fig. 5A). 1: IFT172-FL (homozygous): Both alleles contained the eGFP insert at the C-terminus before the stop codon. 2: IFT172-FL (heterozygous): One allele contained the eGFP insert at the C-terminus, the other allele was the unaltered wildtype allele. 3: IFT172 ΔU-box (homozygous): U-box domain deleted in both alleles by the insertion of eGFP after amino acid 1688. 4: IFT172 ΔU-box (heterozygous): U-box domain deleted in one allele; the other allele was the unaltered wildtype allele. Correct knock-in and the absence of indels was confirmed by Sanger sequencing.

### Immunofluorescence staining and microscopy

RPE1 cells were grown on glass coverslips to 70-80 % confluency and serum-starved for 24 h to induce ciliogenesis. Cells were fixed with 3% PFA in PBS for 15 min and permeabilized for 5 min in 0.2% Triton X-100/PBS. Blocking was performed for 30 min with 3% BSA in PBS. A mouse monoclonal antibody against acetylated tubulin (Sigma-Aldrich, Clone 6-11B-1, diluted 1:500) was used to stain for the ciliary axoneme. For protein localization studies, GFP fluorescence was visualized directly. Primary antibodies were diluted in blocking solution and incubated with the cells at room temperature for 2 h or at 4°C overnight. Coverslips were washed three times with PBS and incubated with an anti-mouse secondary antibody conjugated to Cy3 (Jackson Immuno Research Labs, Cat# 715-165-150, diluted 1:500) for 1 h at room temperature. DAPI (40,6-diamidino-2-phenylindole, Sigma-Aldrich) was included with secondary antibodies for DNA staining. Coverlsips were washed three times with PBS and mounted on glass slides in Mowiol (Sigma-Aldrich).

Widefield images were acquired as z-stacks at 0.3 µm intervals using a Zeiss AxioImager M1 microscope equipped with a Zeiss CellObserver equipped with an Apochromat 63×/NA1.4 oil-immersion objective and a CoolSNAP HQ2 camera. Confocal images were acquired as z-stacks at 0.125 µm intervals using a Nikon Ti-2, A1 LFO confocal microscope with a Plan Apo λ 100× NA 1.4 oil objective. Image analysis was performed using Fiji/ImageJ (NIH)^92^ For quantification of ciliation frequencies, maximum intensity projections of z-stacks were generated, and the number of nuclei/cells and cilia were determined using DAPI and acetylated tubulin staining, respectively. For ciliary length measurements, the region of interest was manually defined using the line segment tool. Measurements were obtained from four independent experiments, and 30-60 cells were analyzed per condition and replicate. Graphs were drawn, and statistical analysis was performed using Prism (GraphPad). Data are presented as mean +/- standard error of the mean (SEM) or box-and-whisker plots with horizontal lines showing 25, 50 and 75th percentiles and whiskers extending to minimum and maximum values.

### AlphaFold

All structural models not supported by crystallographic data were predicted using a local installation of AlphaFold v. 2.3.2^43–44^. Visualizations of protein structure was done using PyMOL v. 2.5 (Schrodinger LLC, https://pymol.org).

## Acknowledgments

We thank Jesper L. Karlsen and Rune T. Kidmose for assistance with biocomputing and the Institute of Molecular Biology and Genetics at Aarhus University for computing time. We also thank Michael Knop, Keith Joung, and Benjamin Kleinstiver for reagents and the Danish Molecular Biomedical Imaging Center, University of Southern Denmark, for the use of imaging equipment, supported by Novo Nordisk Foundation (NNF18SA0032928). This work was funded by grants from the Novo Nordisk Foundation (grant numbers NNF15OC00114164 and NNF23OC0085823) and the European Union’s Horizon 2020 research and innovation program Marie Sklodowska-Curie Innovative Training Networks (ITN) grant 861329 to E.L.

## Figures and figure legends

**Figure S1.**
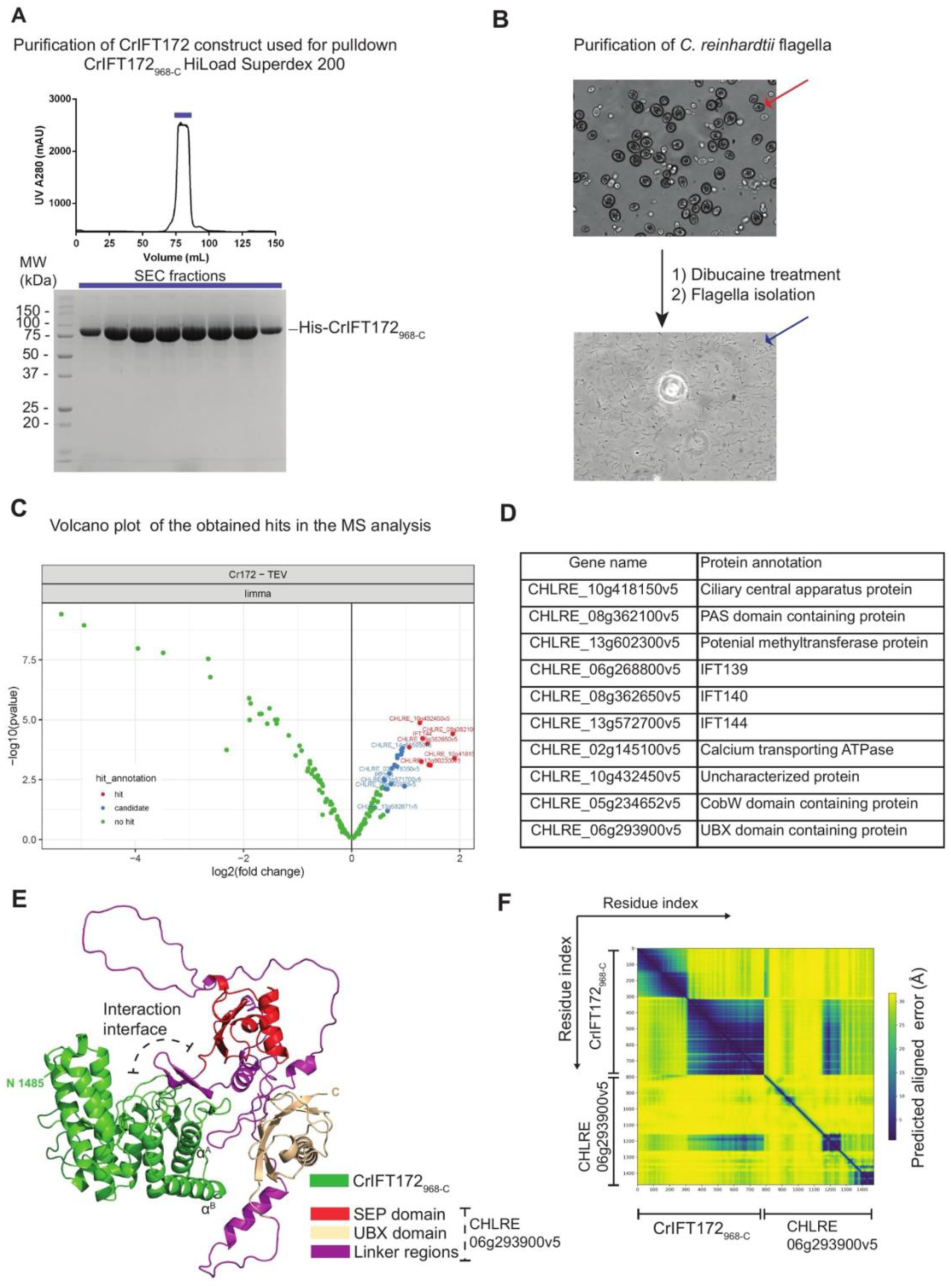
Uncovering interactors of the CrIFT172 C-terminus. **(A)** Size exclusion chromatography (SEC) elution profile for CrIFT172_968-C_ (top). Protein composition of denoted elution fractions analyzed by Coomassie staining after SDS-PAGE separation (bottom). **(B)** *Chlamydomonas reinhardtii* CC1690 cells visualized by light microscopy before flagella isolation (top) and purified flagella fraction after de-flagellation at the same magnification (bottom). **(C)** Volcano plot showing distribution of mass spectrometry (MS) analysis hits for flagellar proteins pulled down by CrIFT172_968-C_ in *C. reinhardtii* CC1690, compared against the tobacco etch virus (TEV) protease control. **(D)** Table of the 10 significant CrIFT172_968-C_ flagellar interactors identified in panel C. Gene name(left) and protein annotations from Uniprot (right) are shown. **(E)** Alphafold predicted structural model for a complex between CrIFT172_968-C_ and the UBX domain containing protein (CHLRE_06g293900v5) identified as an interaction partner (Fig. S1C-D). A linker region that lies between the UBX and SEP domains is predicted to form contacts with IFT172. The IFT-A binding helices described in Fig. 1F are denoted as α^A^ and α^B^. **(F)** PAE plot for the AlphaFold predicted structure shown in panel E.

**Figure S2.**
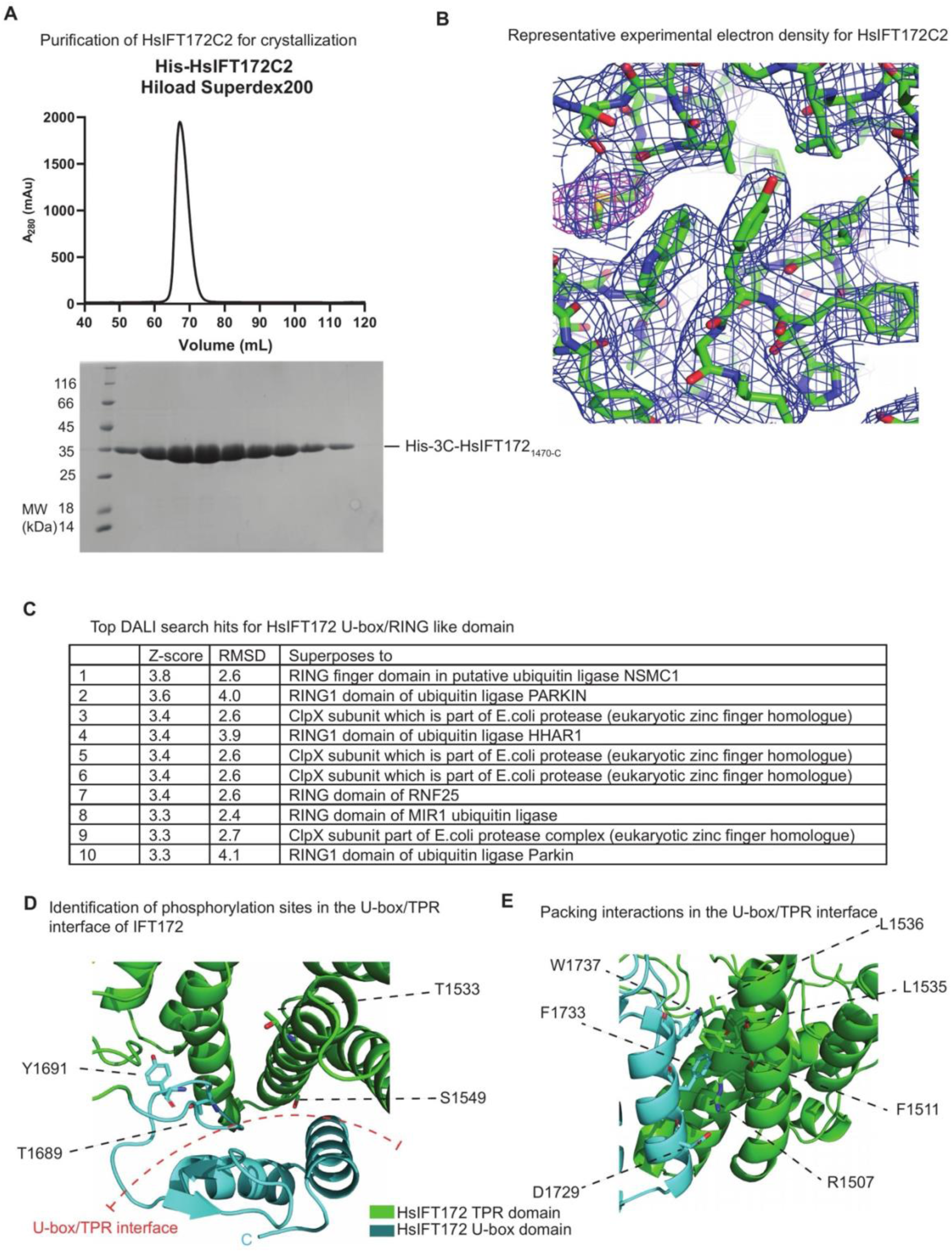
Purification and structure determination of HsIFT172 C-terminal domain. **(A)** SEC elution profile for HsIFT172C2 shown on top. The protein composition of the denoted elution fractions was analyzed by Coomassie staining on an SDS-PAGE (bottom). **(B)** Representative electron density map for HsIFT172C2 crystals. The MR-SAD map is shown as a blue mesh contoured at 1σ and SeMet anomalous density is shown as a magenta mesh contoured at 5σ. **(C)** Top 10 hits obtained from a search of structural homologs for HsIFT172C3 against the PDB database using the DALI server^49^. Z-score and RMSD of corresponding structural alignment are indicated. **(D)** Phosphorylation sites on HsIFT172C2 identified from curated databases of known phosphorylation sites^76^. The phosphorylation sites on HsIFT172C2 are exclusively present at the TPR/U-box interface, suggesting phosphorylation as a potential mechanism for relieving the structural inhibition of the U-box E2 binding site. **(E)** Representative hydrophobic and polar contacts that allow the IFT172 U-box domain to pack against the TPR helices.

**Figure S3.**
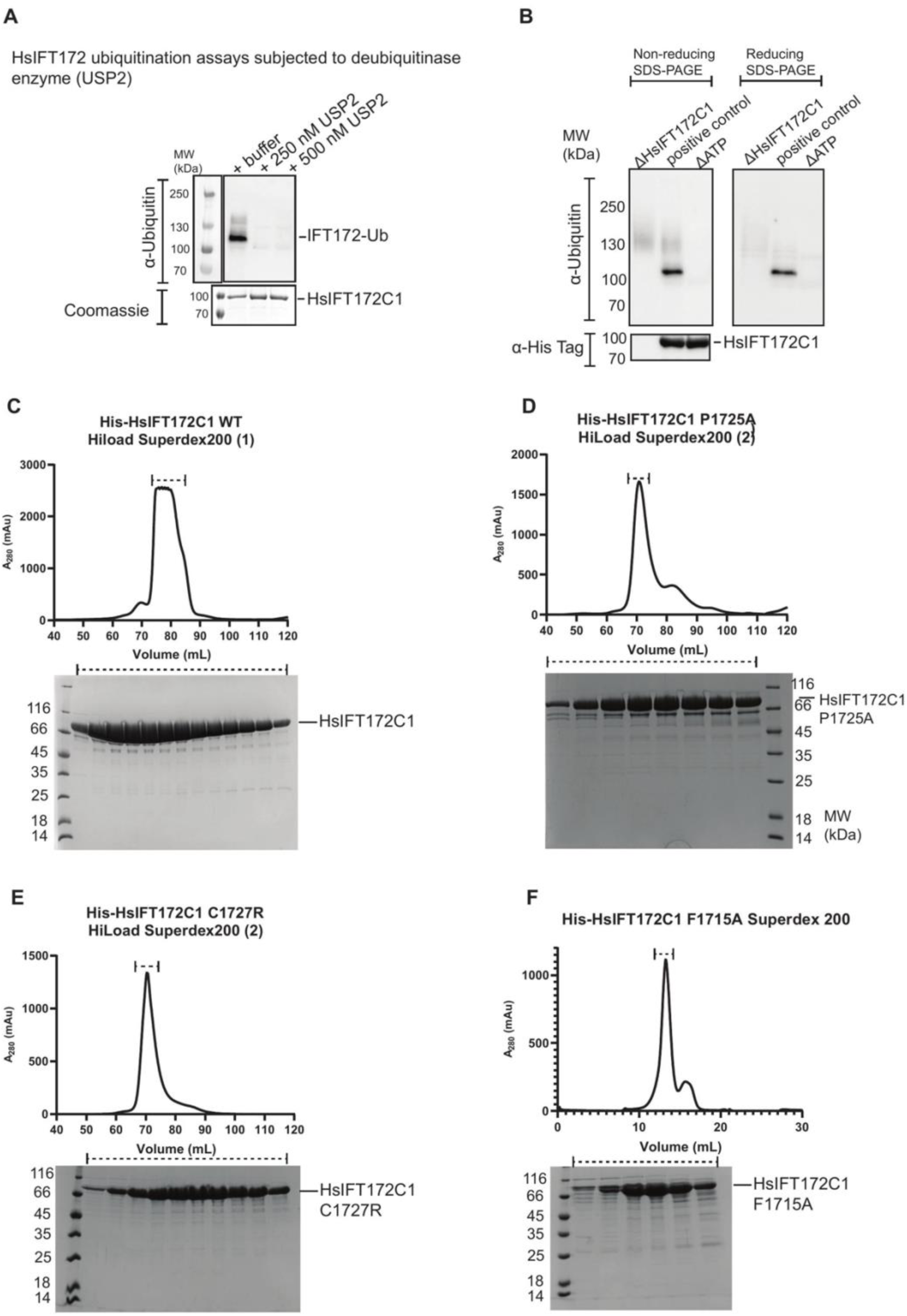
Purification and ubiquitination assays with HsIFT172C constructs. **(A)** *In vitro* ubiquitination reactions performed with HsIFT172C1, followed by incubation with the specified concentrations of the deubiquitinase enzyme USP2 (R&D Systems # E-504-050) or buffer control. Reactions were visualized by immunostaining with (top) anti-Ubiquitin antibody and (bottom) Coomassie staining. **(B)** Western blot analysis of *in vitro* ubiquitination reactions containing HsIFT172C1 after separation on a non-reducing SDS-PAGE (left) and reducing SDS-PAGE gel (right). Reactions were visualized by immunostaining with (top) anti-Ubiquitin antibody and (bottom) anti-His tag antibody. Δ indicates reaction components that were omitted in the specific reaction. **(C-F)** SEC elution profiles (top) and protein composition of the denoted elution fractions analyzed by Coomassie staining post separation on SDS-PAGE (bottom) for: **(C)** HsIFT172C1 WT. **(D)** HsIFT172C1 P1725A. **(E)** HsIFT172C1 C1727R. **(F)** HsIFT172C1 F1715A. (1) and (2) denotes two different HiLoad Superdex 200 columns used to run the samples.

**Figure S4.**
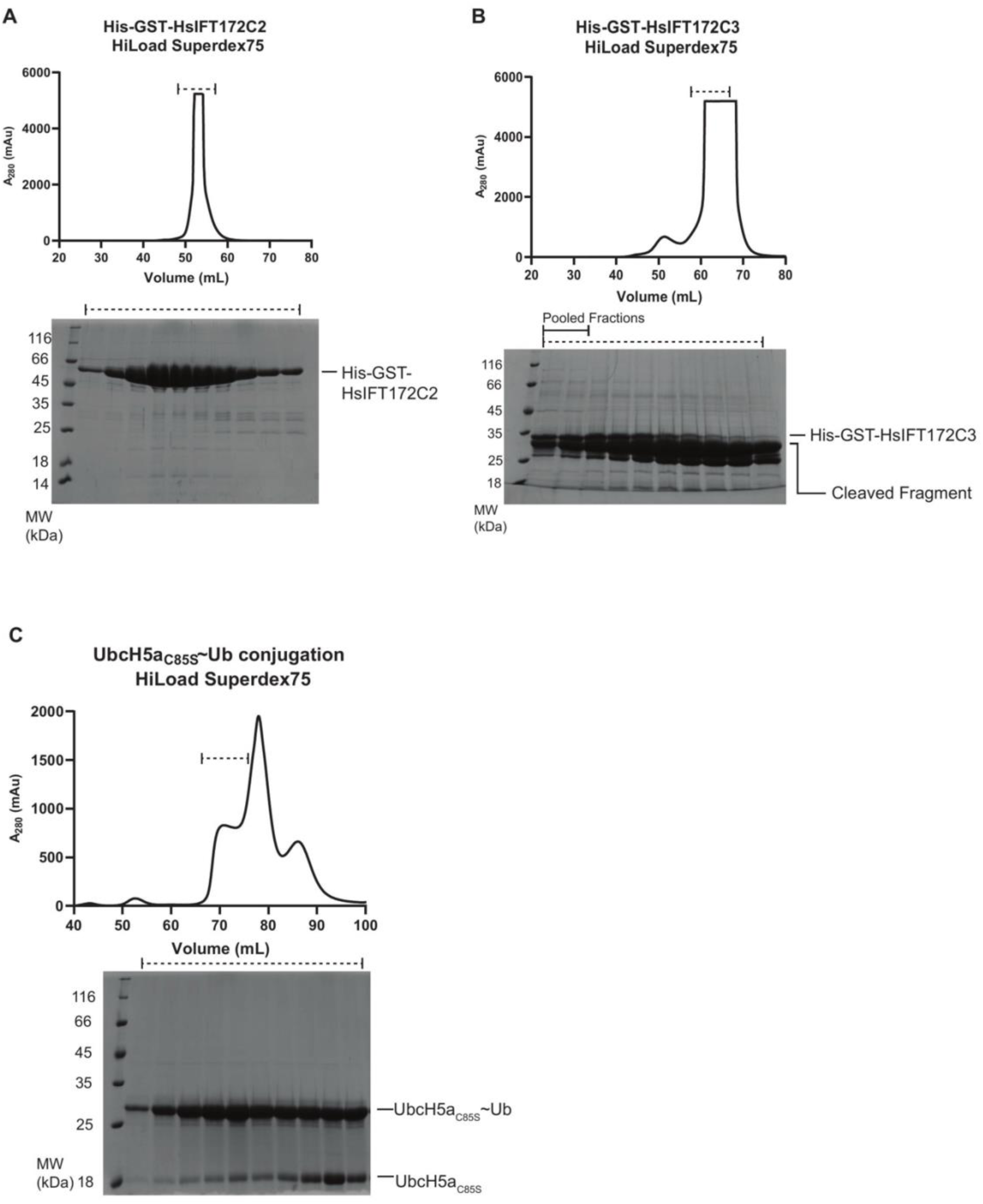
Purification of GST-tagged HsIFT172 constructs and of the E2∼Ub conjugate. **(A)** SEC elution profile (top) and protein composition of the denoted elution fractions analyzed by Coomassie staining after separation on SDS-PAGE (bottom) for His-GST-TEV-HsIFT172C2. **(B)** SEC elution profile (top) and protein composition of the denoted elution fractions (bottom) for His-GST-TEV-HsIFT172C3. A significant amount of proteolytically cleaved protein fragments was observed upon His-GST-TEV-HsIFT172C3 purification. **(C)** Reaction products of an upscaled *in vitro* ubiquitin charging reaction for UbcH5a_C85S_ WT conjugate separated on a SEC column (top). The denoted elution fractions were analyzed by Coomassie staining on an SDS-PAGE gel (bottom).

**Figure S5.**
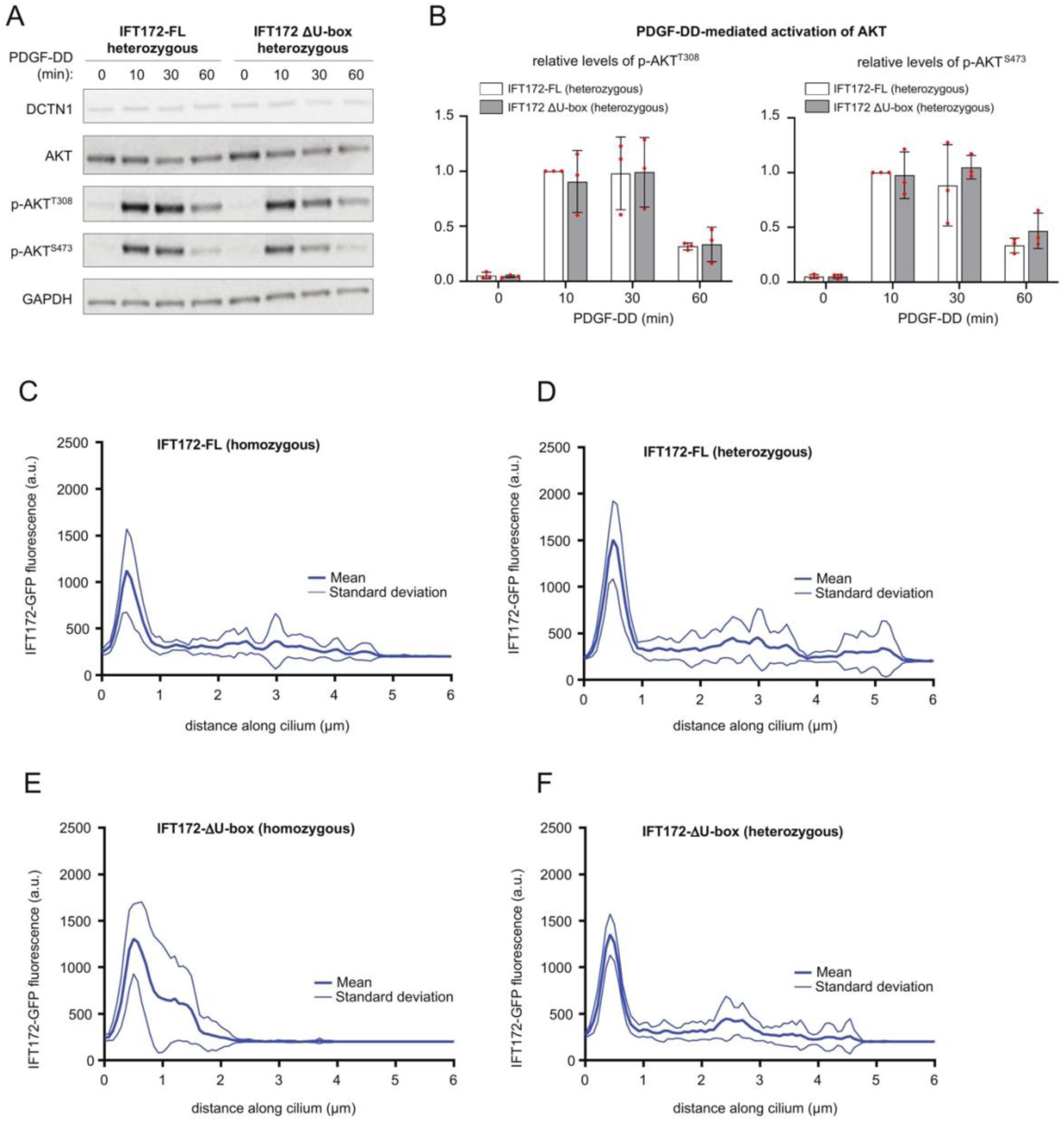
Truncation of the IFT172 U-box domain does not impair PDGF-DD-mediated AKT signaling and ciliary IFT172 localization in RPE1 cells. **(A)** WB analysis of p-AKT levels (p-AKT^T308^ and pAKT^S473^) in IFT172-FL (heterozygous) and IFT172ΔU-box (heterozygous) RPE1 cell lines treated with PDGF-DD ligand for the indicated time points. **(B)** Quantification of p-AKT levels in panel A normalized to DCTN1 and total AKT. Error bars represent standard error of the mean. **(C–F)** Quantification of IFT172-eGFP fluorescence intensity in arbitrary units (a.u.) along the cilium (base to tip) in IFT172-FL (homozygous) **(C)**, IFT172-FL (heterozygous) **(D)**, IFT172ΔU-box (homozygous) **(E)**, and IFT172ΔU-box (heterozygous) **(F)** RPE1 cells. Line scans were performed on n = 10 cilia per cell line. Thick lines, mean; thin lines, standard deviation. The localization pattern of IFT172-eGFP is similar between IFT172-FL and IFT172ΔU-box (heterozygous) cells, whereas IFT172ΔU-box (homozygous) cells display markedly shorter cilia with IFT172 distributed along the full length of the truncated axoneme.

**Figure S6:**
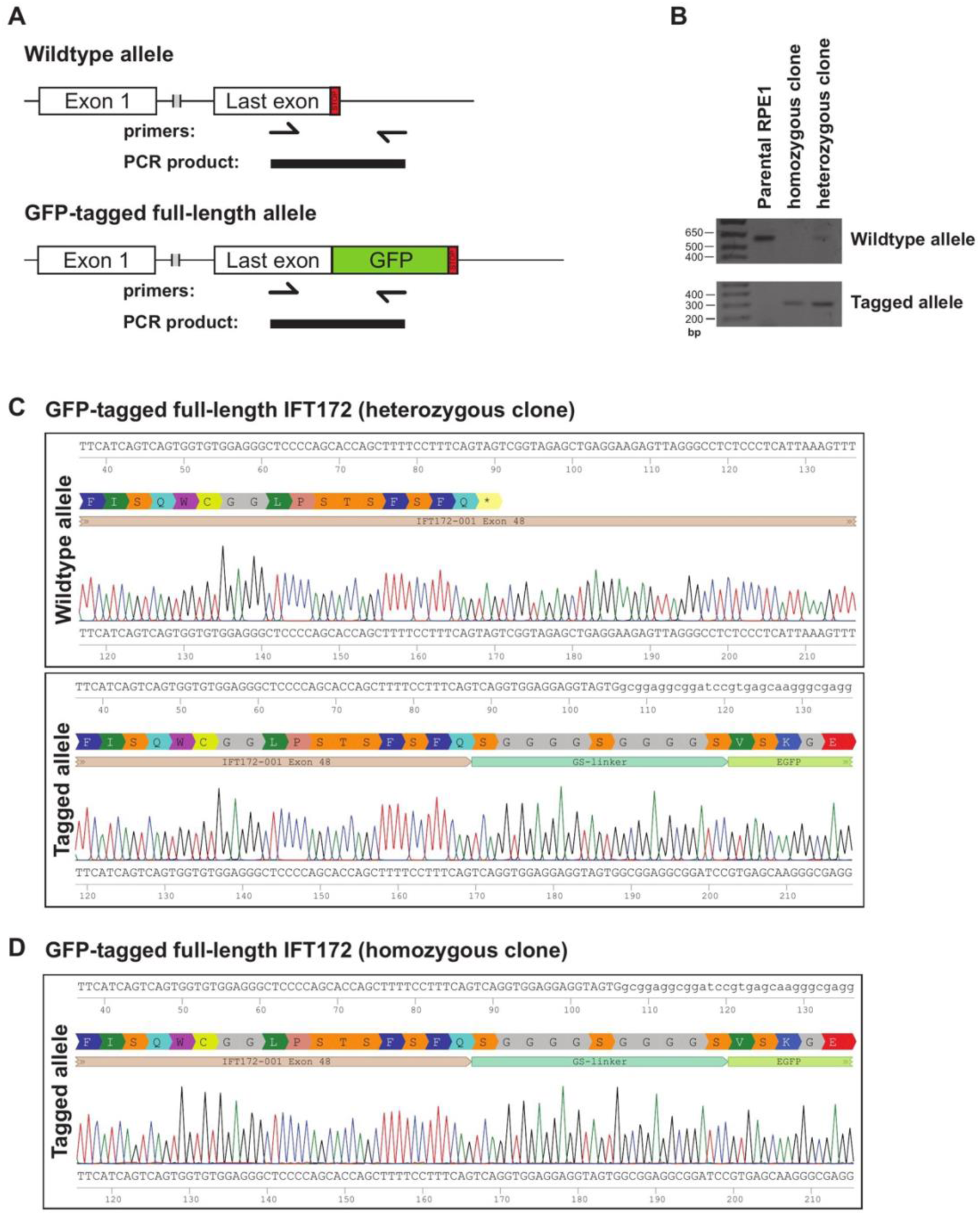
Sequence analysis of GFP-tagged full-length IFT172 clones. **(A)** Tagging and PCR-based screening strategy for cells expressing GFP-tagged full-length IFT172. The GFP tag is inserted into the last exon of IFT172 immediately before the stop codon. Two primer sets are used: the wildtype primer set binds the genomic region flanking the insertion site and amplifies a product spanning the insertion, while the knock-in primer set amplifies only from the successfully tagged allele, as one of its primers binds the inserted GFP sequence. **(B)** Agarose gel electrophoresis of PCR products for the wildtype and tagged alleles amplified from the heterozygous and homozygous cell clones, as well as parental RPE1 cells. **(C)** Sanger sequencing chromatograms of the IFT172 last exon for the heterozygous GFP-tagged clone, showing the wildtype allele (top) and the tagged allele (bottom), confirming correct in-frame insertion of the GFP tag and the absence of indels. **(D)** Sanger sequencing chromatogram of the tagged allele for the homozygous GFP-tagged clone, confirming correct in-frame insertion of the GFP tag in both alleles and the absence of indels.

**Figure S7.**
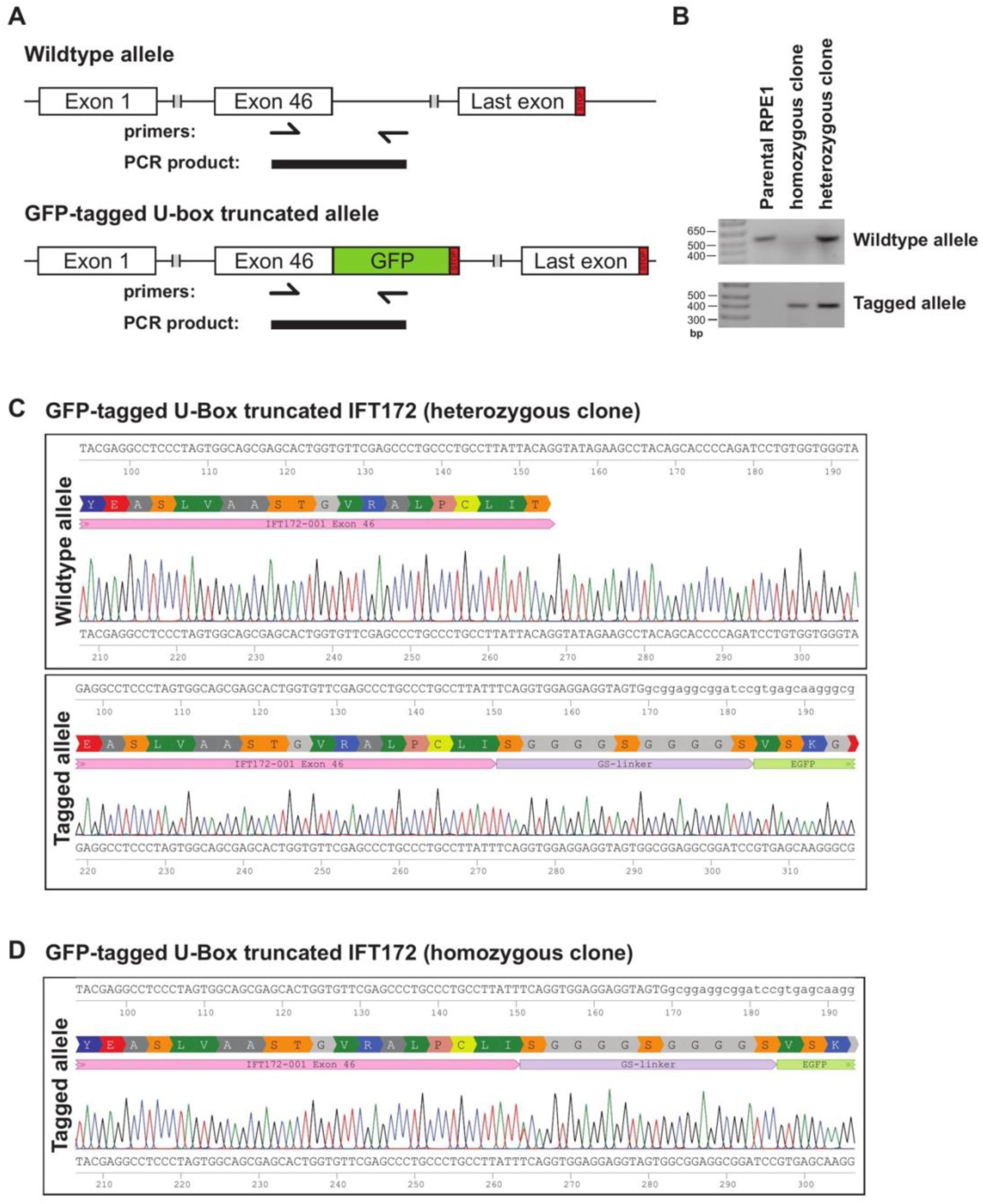
Sequence analysis of GFP-tagged U-box-truncated IFT172 clones. **(A)** Tagging and PCR-based screening strategy for cells expressing GFP-tagged U-box-truncated IFT172. The GFP tag is inserted into exon 46 of IFT172, upstream of the splice donor site, which introduces a premature stop codon and truncates the protein at the start of the U-box domain. Two primer sets are used: the wildtype primer set binds the genomic region flanking the insertion site and amplifies a product spanning the insertion, while the knock-in primer set amplifies only from the successfully tagged allele, as one of its primers binds the inserted GFP sequence. **(B)** Agarose gel electrophoresis of PCR products for the wildtype and tagged alleles amplified from the heterozygous and homozygous cell clones, as well as parental RPE1 cells. **(C)** Sanger sequencing chromatograms of the IFT172 exon 46 region for the heterozygous U-box-truncated clone, showing the wildtype allele (top) and the tagged allele (bottom), confirming correct in-frame insertion of the GFP tag and the absence of indels. **(D)** Sanger sequencing chromatogram of the tagged allele for the homozygous U-box-truncated clone, confirming correct in-frame insertion of the GFP tag in both alleles and the absence of indels.

**Figure S8.**
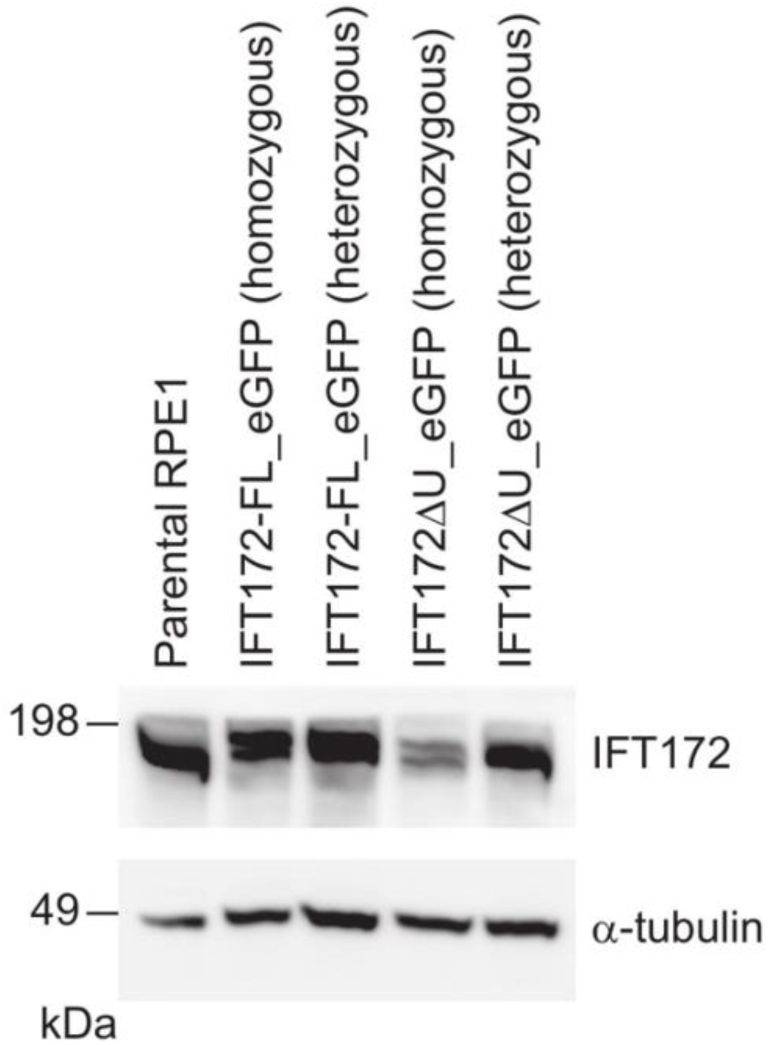
Immunoblot analysis of IFT172 expression in the engineered RPE1 cell lines. Whole-cell lysates from parental RPE1 cells and the four engineered cell lines (IFT172-FL_eGFP homozygous and heterozygous; IFT172ΔU-box_eGFP homozygous and heterozygous) were analyzed by immunoblotting with an anti-IFT172 antibody. α-tubulin was used as a loading control. Expression of the tagged full-length and U-box-truncated IFT172 proteins is confirmed in both homozygous and heterozygous clones. Reduced steady-state levels of IFT172 are observed in the IFT172ΔU-box_eGFP (homozygous) clone compared to the IFT172-FL_eGFP clones, consistent with compromised protein stability upon deletion of the U-box domain.

## References

1. Satir, P. & Christensen, S. T. Overview of Structure and Function of Mammalian Cilia. Annu. Rev. Physiol. 69, 377–400 (2007).

2. Wan, K. Y. Coordination of eukaryotic cilia and flagella. Essays Biochem. 62, 829–838 (2018).

3. Mill, P., Christensen, S. T. & Pedersen, L. B. Primary cilia as dynamic and diverse signalling hubs in development and disease. Nat. Rev. Genet. 24, 421–441 (2023).

4. Reiter, J. F. & Leroux, M. R. Genes and molecular pathways underpinning ciliopathies. Nat. Rev. Mol. Cell Biol. 18, 533–547 (2017).

5. Focşa, I. O., Budişteanu, M. & Bălgrădean, M. Clinical and genetic heterogeneity of primary ciliopathies (Review). Int. J. Mol. Med. 48, 176 (2021).

6. Rosenbaum, J. L. & Witman, G. B. Intraflagellar transport. Nat. Rev. Mol. Cell Biol. 3, 813–825 (2002).

7. Kozminski, K. G., Johnson, K. A., Forscher, P. & Rosenbaum, J. L. A motility in the eukaryotic flagellum unrelated to flagellar beating. Proc. Natl. Acad. Sci. U. S. A. 90, 5519–5523 (1993).

8. Kozminski, K. G., Beech, P. L. & Rosenbaum, J. L. The Chlamydomonas kinesin-like protein FLA10 is involved in motility associated with the flagellar membrane. J. Cell Biol. 131, 1517–1527 (1995).

9. Bhogaraju, S., Weber, K., Engel, B. D., Lechtreck, K. & Lorentzen, E. Getting tubulin to the tip of the cilium: One IFT train, many different tubulin cargo-binding sites? BioEssays 36, 463–467 (2014).

10. Lechtreck, K. F. IFT-Cargo Interactions and Protein Transport in Cilia. Trends Biochem. Sci. 40, 765–778 (2015).

11. Long, H. & Huang, K. Transport of Ciliary Membrane Proteins. Front. Cell Dev. Biol. 7, (2020).

12. Mukhopadhyay, S. et al. TULP3 bridges the IFT-A complex and membrane phosphoinositides to promote trafficking of G protein-coupled receptors into primary cilia. Genes Dev. 24, 2180–2193 (2010).

13. Pigino, G. et al. Electron-tomographic analysis of intraflagellar transport particle trains in situ. J. Cell Biol. 187, 135–148 (2009).

14. van den Hoek, H., et al. In situ architecture of the ciliary base reveals the stepwise assembly of intraflagellar transport trains. Science 377, 543–548 (2022).

15. Taschner, M. & Lorentzen, E. The Intraflagellar Transport Machinery. Cold Spring Harb. Perspect. Biol. a028092 (2016) doi:10.1101/cshperspect.a028092.

16. Wingfield, J. L. et al. IFT trains in different stages of assembly queue at the ciliary base for consecutive release into the cilium. eLife 6, e26609 (2017).

17. Pedersen, L. B., Geimer, S. & Rosenbaum, J. L. Dissecting the Molecular Mechanisms of Intraflagellar Transport in Chlamydomonas. Curr. Biol. 16, 450–459 (2006).

18. Jordan, M. A., Diener, D. R., Stepanek, L. & Pigino, G. The cryo-EM structure of intraflagellar transport trains reveals how dynein is inactivated to ensure unidirectional anterograde movement in cilia. Nat. Cell Biol. 20, 1250–1255 (2018).

19. Pazour, G. J., Dickert, B. L. & Witman, G. B. The DHC1b (DHC2) isoform of cytoplasmic dynein is required for flagellar assembly. J. Cell Biol. 144, 473–481 (1999).

20. Porter, M. E., Bower, R., Knott, J. A., Byrd, P. & Dentler, W. Cytoplasmic dynein heavy chain 1b is required for flagellar assembly in Chlamydomonas. Mol. Biol. Cell 10, 693–712 (1999).

21. Eguether, T. et al. IFT27 Links the BBSome to IFT for Maintenance of the Ciliary Signaling Compartment. Dev. Cell 31, 279–290 (2014).

22. Nachury, M. V. et al. A Core Complex of BBS Proteins Cooperates with the GTPase Rab8 to Promote Ciliary Membrane Biogenesis. Cell 129, 1201–1213 (2007).

23. Komander, D. & Rape, M. The Ubiquitin Code. Annu. Rev. Biochem. 81, 203–229 (2012).

24. Hossain, D. & Tsang, W. Y. The role of ubiquitination in the regulation of primary cilia assembly and disassembly. Semin. Cell Dev. Biol. 93, 145–152 (2019).

25. Lv, B., Stuck, M. W., Desai, P. B., Cabrera, O. A. & Pazour, G. J. E3 ubiquitin ligase Wwp1 regulates ciliary dynamics of the Hedgehog receptor Smoothened. J. Cell Biol. 220, e202010177 (2021).

26. Shinde, S. R., Nager, A. R. & Nachury, M. V. Ubiquitin chains earmark GPCRs for BBSome-mediated removal from cilia. J. Cell Biol. 219, e202003020 (2020).

27. Huang, K., Diener, D. R. & Rosenbaum, J. L. The ubiquitin conjugation system is involved in the disassembly of cilia and flagella. J. Cell Biol. 186, 601–613 (2009).

28. Wang, Q., Peng, Z., Long, H., Deng, X. & Huang, K. Polyubiquitylation of α-tubulin at K304 is required for flagellar disassembly in Chlamydomonas. J. Cell Sci. 132, jcs229047 (2019).

29. Desai, P. B., Stuck, M. W., Lv, B. & Pazour, G. J. Ubiquitin links smoothened to intraflagellar transport to regulate Hedgehog signaling. J. Cell Biol. 219, e201912104 (2020).

30. Kim, J. et al. The role of ciliary trafficking in Hedgehog receptor signaling. Sci. Signal. 8, ra55 (2015).

31. Shinde, S. R. et al. The ancestral ESCRT protein TOM1L2 selects ubiquitinated cargoes for retrieval from cilia. Dev. Cell 58, 677–693.e9 (2023).

32. Aslanyan, M. G. et al. A targeted multi-proteomics approach generates a blueprint of the ciliary ubiquitinome. Front. Cell Dev. Biol. 11, (2023).

33. Chiuso, F. et al. Ubiquitylation of BBSome is required for ciliary assembly and signaling. EMBO Rep. 24, e55571 (2023).

34. Taschner, M. et al. Intraflagellar transport proteins 172, 80, 57, 54, 38, and 20 form a stable tubulin-binding IFT-B2 complex. EMBO J. 35, 773–790 (2016).

35. Petriman, N. A. et al. Biochemically validated structural model of the 15-subunit intraflagellar transport complex IFT-B. EMBO J. 41, e112440 (2022).

36. Halbritter, J. et al. Defects in the IFT-B Component IFT172 Cause Jeune and Mainzer-Saldino Syndromes in Humans. Am. J. Hum. Genet. 93, 915–925 (2013).

37. Bujakowska, K. M. et al. Mutations in IFT172 cause isolated retinal degeneration and Bardet–Biedl syndrome. Hum. Mol. Genet. 24, 230–242 (2015).

38. Lucas-Herald, A. K. et al. A Case of Functional Growth Hormone Deficiency and Early Growth Retardation in a Child With IFT172 Mutations. J. Clin. Endocrinol. Metab. 100, 1221–1224 (2015).

39. Huangfu, D. et al. Hedgehog signalling in the mouse requires intraflagellar transport proteins. Nature 426, 83–87 (2003).

40. Pedersen, L. B. et al. Chlamydomonas IFT172 Is Encoded by FLA11, Interacts with CrEB1, and Regulates IFT at the Flagellar Tip. Curr. Biol. 15, 262–266 (2005).

41. Tsao, C.-C. & Gorovsky, M. A. Different Effects of Tetrahymena IFT172 Domains on Anterograde and Retrograde Intraflagellar Transport. Mol. Biol. Cell 19, 1450–1461 (2008).

42. Williamson, S. M., Silva, D. A., Richey, E. & Qin, H. Probing the role of IFT particle complex A and B in flagellar entry and exit of IFT-dynein in Chlamydomonas. Protoplasma 249, 851–856 (2012).

43. Jumper, J. et al. Highly accurate protein structure prediction with AlphaFold. Nature 596, 583–589 (2021).

44. Evans, R. et al. Protein complex prediction with AlphaFold-Multimer. 2021.10.04.463034 Preprint at 10.1101/2021.10.04.463034 (2022).

45. Jiang, M. et al. Human IFT-A complex structures provide molecular insights into ciliary transport. Cell Res. 33, 288–298 (2023).

46. Lacey, S. E., Foster, H. E. & Pigino, G. The molecular structure of IFT-A and IFT-B in anterograde intraflagellar transport trains. Nat. Struct. Mol. Biol. 30, 584–593 (2023).

47. Lacey, S. E., Graziadei, A. & Pigino, G. Extensive structural rearrangement of intraflagellar transport trains underpins bidirectional cargo transport. Cell 187, 4621–4636.e18 (2024).

48. Kloppsteck, P., Ewens, C. A., Förster, A., Zhang, X. & Freemont, P. S. Regulation of p97 in the ubiquitin–proteasome system by the UBX protein-family. Biochim. Biophys. Acta BBA - Mol. Cell Res. 1823, 125–129 (2012).

49. Holm, L. Using Dali for Protein Structure Comparison. in Structural Bioinformatics: Methods and Protocols (ed. Gáspári, Z.) 29–42 (Springer US, New York, NY, 2020). doi:10.1007/978-1-0716-0270-6_3.

50. Zheng, N. & Shabek, N. Ubiquitin Ligases: Structure, Function, and Regulation. Annu. Rev. Biochem. 86, 129–157 (2017).

51. Ohi, M. D., Vander Kooi, C. W., Rosenberg, J. A., Chazin, W. J. & Gould, K. L. Structural insights into the U-box, a domain associated with multi-ubiquitination. Nat. Struct. Mol. Biol. 10, 250–255 (2003).

52. Aravind, L. & Koonin, E. V. The U box is a modified RING finger — a common domain in ubiquitination. Curr. Biol. 10, R132–R134 (2000).

53. Hatakeyama, S., Yada, M., Matsumoto, M., Ishida, N. & Nakayama, K.-I. U Box Proteins as a New Family of Ubiquitin-Protein Ligases *. J. Biol. Chem. 276, 33111–33120 (2001).

54. Özkan, E., Yu, H. & Deisenhofer, J. Mechanistic insight into the allosteric activation of a ubiquitin-conjugating enzyme by RING-type ubiquitin ligases. Proc. Natl. Acad. Sci. 102, 18890–18895 (2005).

55. Dou, H., Buetow, L., Sibbet, G. J., Cameron, K. & Huang, D. T. BIRC7–E2 ubiquitin conjugate structure reveals the mechanism of ubiquitin transfer by a RING dimer. Nat. Struct. Mol. Biol. 19, 876–883 (2012).

56. Plechanovová, A., Jaffray, E. G., Tatham, M. H., Naismith, J. H. & Hay, R. T. Structure of a RING E3 ligase and ubiquitin-loaded E2 primed for catalysis. Nature 489, 115–120 (2012).

57. Soss, S. E., Klevit, R. E. & Chazin, W. J. Activation of UbcH5c∼Ub is the result of a shift in interdomain motions of the conjugate bound to U-box E3 ligase E4B. Biochemistry 52, 2991–2999 (2013).

58. Buetow, L. & Huang, D. T. Structural insights into the catalysis and regulation of E3 ubiquitin ligases. Nat. Rev. Mol. Cell Biol. 17, 626–642 (2016).

59. Dou, H. et al. Structural basis for autoinhibition and phosphorylation-dependent activation of c-Cbl. Nat. Struct. Mol. Biol. 19, 184–192 (2012).

60. Trempe, J.-F. et al. Structure of parkin reveals mechanisms for ubiquitin ligase activation. Science 340, 1451–1455 (2013).

61. Moura, T. R. de et al. Prp19/Pso4 Is an Autoinhibited Ubiquitin Ligase Activated by Stepwise Assembly of Three Splicing Factors. Mol. Cell 69, 979–992.e6 (2018).

62. de Bie, P. & Ciechanover, A. Ubiquitination of E3 ligases: self-regulation of the ubiquitin system via proteolytic and non-proteolytic mechanisms. Cell Death Differ. 18, 1393–1402 (2011).

63. Hoeller, D. et al. E3-Independent Monoubiquitination of Ubiquitin-Binding Proteins. Mol. Cell 26, 891–898 (2007).

64. Garcia-Barcena, C., Osinalde, N., Ramirez, J. & Mayor, U. How to Inactivate Human Ubiquitin E3 Ligases by Mutation. Front. Cell Dev. Biol. 8, (2020).

65. Zheng, N., Wang, P., Jeffrey, P. D. & Pavletich, N. P. Structure of a c-Cbl–UbcH7 Complex: RING Domain Function in Ubiquitin-Protein Ligases. Cell 102, 533–539 (2000).

66. Middleton, A. J., Budhidarmo, R. & Day, C. L. Chapter Ten - Use of E2∼Ubiquitin Conjugates for the Characterization of Ubiquitin Transfer by RING E3 Ligases Such as the Inhibitor of Apoptosis Proteins. in Methods in Enzymology (eds. Ashkenazi, A., Wells, J. A. & Yuan, J.) vol. 545 243–263 (Academic Press, 2014).

67. Gundogdu, M. & Walden, H. Structural basis of generic versus specific E2–RING E3 interactions in protein ubiquitination. Protein Sci. 28, 1758–1770 (2019).

68. Levin, I. et al. Identification of an unconventional E3 binding surface on the UbcH5 ∼ Ub conjugate recognized by a pathogenic bacterial E3 ligase. Proc. Natl. Acad. Sci. 107, 2848–2853 (2010).

69. Xu, Z. et al. Interactions between the quality control ubiquitin ligase CHIP and ubiquitin conjugating enzymes. BMC Struct. Biol. 8, 26 (2008).

70. Pruski, M. et al. Roles for IFT172 and Primary Cilia in Cell Migration, Cell Division, and Neocortex Development. Front. Cell Dev. Biol. 7, (2019).

71. Clement, C. A. et al. TGF-β Signaling Is Associated with Endocytosis at the Pocket Region of the Primary Cilium. Cell Rep. 3, 1806–1814 (2013).

72. Doganli, C. et al. TAK1 operates at the primary cilium in non-canonical TGFB/BMP signaling to control heart development. 2024.05.06.592628 Preprint at 10.1101/2024.05.06.592628 (2024).

73. Nielsen, B. S. et al. PDGFRβ and oncogenic mutant PDGFRα D842V promote disassembly of primary cilia through a PLCγ- and AURKA-dependent mechanism. J. Cell Sci. 128, 3543–3549 (2015).

74. Wang, Q. et al. Membrane association and remodeling by intraflagellar transport protein IFT172. Nat. Commun. 9, 4684 (2018).

75. Kassenbrock, C. K. & Anderson, S. M. Regulation of Ubiquitin Protein Ligase Activity in c-Cbl by Phosphorylation-induced Conformational Change and Constitutive Activation by Tyrosine to Glutamate Point Mutations*. J. Biol. Chem. 279, 28017–28027 (2004).

76. Hornbeck, P. V. et al. PhosphoSitePlus, 2014: mutations, PTMs and recalibrations. Nucleic Acids Res. 43, D512–D520 (2015).

77. Bomar, M. G., Pai, M.-T., Tzeng, S.-R., Li, S. S.-C. & Zhou, P. Structure of the ubiquitin-binding zinc finger domain of human DNA Y-polymerase η. EMBO Rep. 8, 247–251 (2007).

78. Guo, X. & Wang, X.-F. Signaling cross-talk between TGF-β/BMP and other pathways. Cell Res. 19, 71–88 (2009).

79. Heldin, C.-H. & Moustakas, A. Signaling Receptors for TGF-β Family Members. Cold Spring Harb. Perspect. Biol. 8, a022053 (2016).

80. Zhang, Y. E. Non-Smad pathways in TGF-β signaling. Cell Res. 19, 128–139 (2009).

81. Gibson, D. G. et al. Enzymatic assembly of DNA molecules up to several hundred kilobases. Nat. Methods 6, 343–345 (2009).

82. Craige, B., Brown, J. M. & Witman, G. B. Isolation of Chlamydomonas Flagella. Curr. Protoc. Cell Biol. 59, 3.41.1–3.41.9 (2013).

83. Kabsch, W. XDS. Acta Crystallogr. D Biol. Crystallogr. 66, 125–132 (2010).

84. Winn, M. D. et al. Overview of the CCP4 suite and current developments. Acta Crystallogr. D Biol. Crystallogr. 67, 235–242 (2011).

85. McCoy, A. J. et al. Phaser crystallographic software. J. Appl. Crystallogr. 40, 658–674 (2007).

86. Liebschner, D. et al. Macromolecular structure determination using X-rays, neutrons and electrons: recent developments in Phenix. Acta Crystallogr. Sect. Struct. Biol. 75, 861–877 (2019).

87. Hughes, C. S. et al. Single-pot, solid-phase-enhanced sample preparation for proteomics experiments. Nat. Protoc. 14, 68–85 (2019).

88. Kuhns, S. et al. Endogenous Tagging of Ciliary Genes in Human RPE1 Cells for Live-Cell Imaging. Methods Mol Biol 2725, 147–166 (2024).

89. Fueller, J. et al. CRISPR-Cas12a-assisted PCR tagging of mammalian genes. J Cell Biol 219, e201910210 (2020).

90. Kleinstiver, B. P. et al. Engineered CRISPR-Cas12a variants with increased activities and improved targeting ranges for gene, epigenetic and base editing. Nat Biotechnol 37, 276–282 (2019).

91. Riesenberg, S. et al. Simultaneous precise editing of multiple genes in human cells. Nucleic Acids Res 47, e116 (2019).

92. Schindelin, J., et al. Fiji: an open-source platform for biological-image analysis. Nat Methods 9, 676–682 (2012).

93. Emsley, P., Lohkamp, B., Scott, W. G. & Cowtan, K. Features and development of Coot. Acta Crystallogr D Biol Crystallogr 66, 486–501 (2010).

94. Raman, M., et al. Systematic proteomics of the VCP–UBXD adaptor network identifies a role for UBXN10 in regulating ciliogenesis. Nat. Cell Biol. 17, 1356–1369 (2015).

95. Jiang, M., et al. Human IFT-A complex structures provide molecular insights into ciliary transport. Cell Research 33:288–298 (2023).

96. Walia, V., et al. Akt regulates a Rab11-effector switch required for ciliogenesis. Developmental Cell 50:229–246 (2019).

97. Shim, M. S. et al. Primary cilia and the reciprocal activation of AKT and SMAD2/3 regulate stretch-induced autophagy in trabecular meshwork cells. Proc. Natl. Acad. Sci. U.S.A. 118:e2021942118 (2021).

